# Scalable biophysical constraints link steady-state and kinetic metabolic modeling

**DOI:** 10.64898/2026.07.03.736321

**Authors:** Ilias Toumpe, Daniel R. Weilandt, Bharath Narayanan, Georgios Fengos, Vassily Hatzimanikatis, Ljubisa Miskovic

## Abstract

Rational metabolic engineering relies on models that connect molecular mechanisms to cellular behavior. Constraint-based metabolic models integrate stoichiometric, thermodynamic, and omics-derived information, but they do not generally constrain whether flux magnitudes are compatible with the corresponding metabolite pool sizes. Metabolite turnover rates, the ratios of total production or consumption flux to pool size, capture this relationship and quantify how rapidly pools are renewed. Without turnover constraints, such models can admit flux-concentration combinations with implausible pool-renewal timescales despite satisfying mass balance and thermodynamics. We present METEOR-K, a scalable framework that couples steady-state fluxes and metabolite concentrations through turnover-rate constraints and turnover-aware sampling. These constraints encode physiological timescales or experimental turnover estimates, either network-wide or for selected metabolites. We applied METEOR-K to a reduced glycolysis pathway, two *Escherichia coli* networks, one anaerobic and one aerobic, and near-genome-scale ovarian cancer metabolism. Across these systems, METEOR-K narrowed the admissible steady-state space, reduced flux-concentration uncertainty, and shifted kinetic ensembles toward faster local dynamics and physiologically compatible responses. Nonlinear simulations yielded *E. coli* growth behavior consistent with experimental observations and supported perturbation recovery in the ovarian cancer model. In the aerobic *E. coli* core model, literature-derived turnover intervals for 18 metabolites narrowed flux and concentration ranges across the network, increased the fraction of locally stable models from 26.8% to 73.5%, and shortened median relaxation times by over four orders of magnitude relative to the unconstrained reference. METEOR-K provides turnover-consistent steady states for constraint-based analyses and physiologically grounded reference states for kinetic modeling and model-guided metabolic engineering.

## Introduction

Rational metabolic engineering depends on accurate and reliable models that connect molecular mechanisms to cellular behavior and predict the effects of genetic, environmental, and process interventions. Developing such models remains challenging because cellular metabolism is complex and exhibits various nonlinear dynamic phenomena, including negative feedback and feedforward loops ^1,2^, oscillations ^3–5^, and multiple steady states ^6–9^. Although models used for rational design need not reproduce all these phenomena explicitly, the steady states underlying their predictions should be compatible with physiological metabolic timescales. Otherwise, engineering targets and intervention responses may be predicted from biologically unrealistic operating states. At the genome scale, this creates a need to combine scalable characterization of feasible steady states with an assessment of their dynamic plausibility.

Constraint-based modeling has been central to the scalable characterization of steady states because it enables large-scale analysis of steady-state metabolic fluxes under physicochemical and physiological constraints. These methods identify flux states consistent with mass balance ^10^, thermodynamics ^11–14^, resource allocation ^15,16^, and multi-omics data integration ^17–20^. However, even with these extensions, the feasible solution space often remains large, and not all admissible flux states correspond to physiologically plausible metabolic behavior.

A key limitation is that steady-state formulations generally do not constrain how flux magnitudes scale with metabolite pool sizes ^21,22^. For example, thermodynamics-based formulations ^11–14^ incorporate metabolite concentrations via Gibbs-energy constraints, but these primarily restrict reaction directionality and feasible concentration ranges. They do not determine whether a flux magnitude is kinetically plausible for the corresponding metabolite pool size. Thus, flux distributions that satisfy mass balance, thermodynamic feasibility, and other imposed constraints can still imply unrealistic biochemical dynamics.

The physiological consequences of this missing coupling become apparent as metabolic demand changes. Metabolite concentrations are generally maintained within relatively narrow physiological ranges, whereas metabolic fluxes must adjust more broadly to support growth and environmental responses. Cells therefore cannot simply increase metabolite pools in direct proportion to flux. Instead, higher flux through constrained pools requires faster metabolite renewal, linking flux-concentration relationships to metabolic timescales ^23^. In growing cells, a steady-state flux-concentration pair that implies metabolite renewal timescales much slower than the growth process is dynamically inconsistent with the physiology it is intended to represent, even if it satisfies mass balance and thermodynamic feasibility.

Metabolite turnover rates provide a direct measure of this renewal. For a metabolite, the turnover rate is the ratio of the cumulative production or consumption rate to its concentration, and the corresponding turnover time is its inverse. This ratio links a steady-state flux-concentration pair to an implied pool-renewal timescale. In the simplest first-order case, where the flux, *v*, is a linear function of the metabolite concentration, *x*, (*v* = k · *x*), the ratio *v*/*x* is the kinetic constant *k*. Thus, a flux-concentration pair already encodes a characteristic timescale without requiring a full kinetic model (Figure 1).

**Figure 1.**
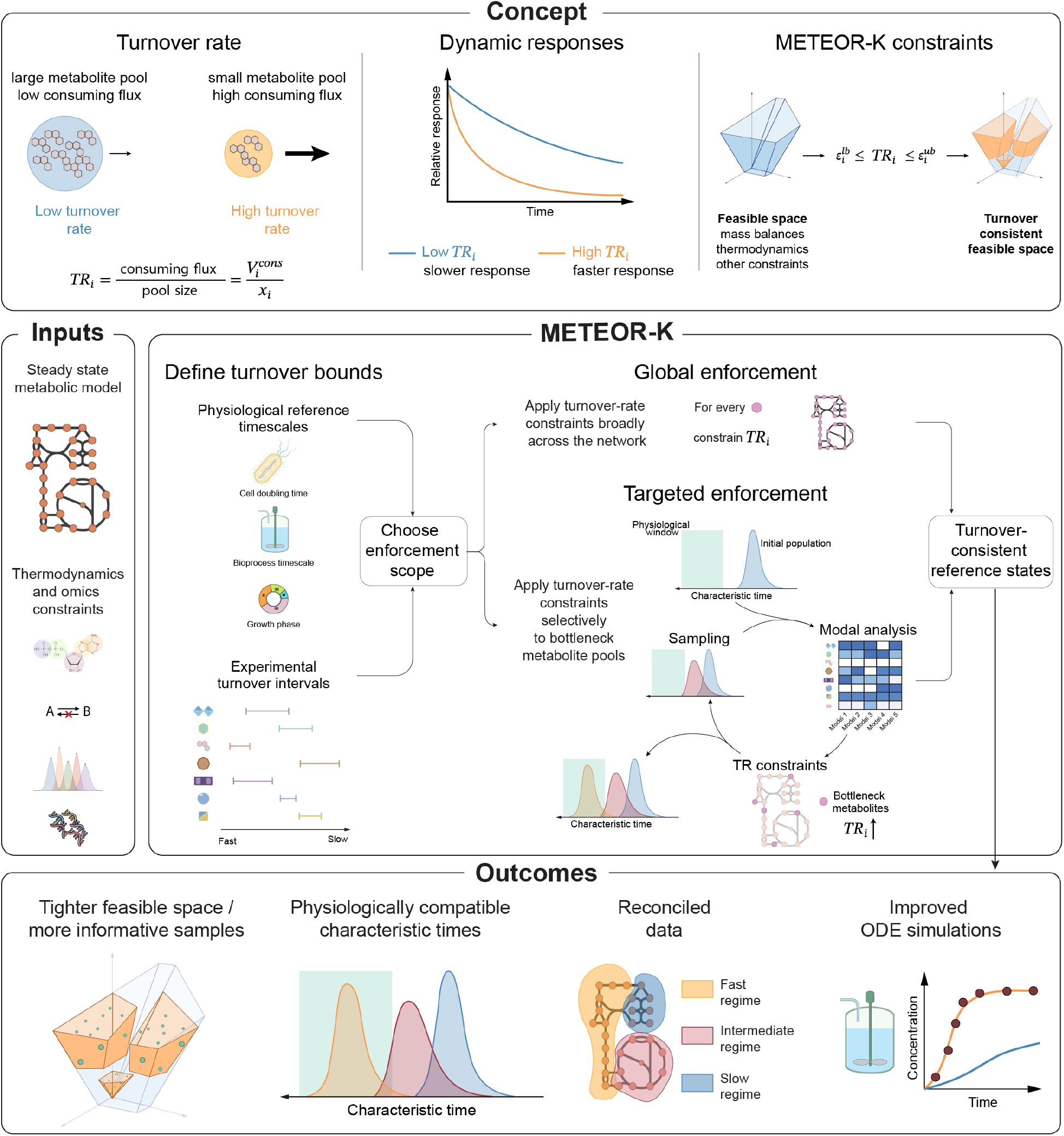
Turnover rate (TR) constraints and the METEOR-K workflow. The flux-concentration ratio defines an implied turnover rate for each metabolite pool, linking pool size and consuming flux to local dynamic response timescales. METEOR-K couples flux and concentration variables by enforcing bounds on metabolite TRs, thereby restricting the feasible space to turnover-consistent metabolic states. The framework takes as input a steady-state constraint-based metabolic model with flux and concentration ranges under a given physiology, optionally augmented with thermodynamic, exchange, growth, omics, and pathway-level timescale constraints. METEOR-K can be applied globally using physiologically relevant reference timescales, such as cell doubling time, bioprocess time, or growth-phase timescales, or in a targeted manner by identifying metabolites associated with slow dynamical modes and imposing TR constraints on those pools. The resulting constrained solution space yields tighter feasible regions, more informative steady-state samples, characteristic times closer to physiological regimes, reconciled data, and improved nonlinear ODE simulations.

A physiologically plausible steady state should also be compatible with realistic local dynamic behavior, which is governed by the Jacobian evaluated at that state. Because Jacobian entries scale with flux-to-concentration ratios (Supplementary Note 1), metabolite turnover rates influence the range of attainable relaxation timescales. However, kinetic sensitivities, regulatory couplings, conserved pools, and network interactions determine how individual dynamic modes are realized within that range. Consequently, turnover-rate constraints provide a dynamic screen: low turnover can indicate steady states likely to support slow local responses, whereas high turnover permits, but does not guarantee, fast dynamics.

To incorporate turnover-based screening for dynamic plausibility into genome-scale constraint-based modeling, we developed METEOR-K (MEtabolite Turnover-rate Enabled Optimization for Relevant Kinetics). This scalable mixed-integer linear programming framework couples steady-state metabolic fluxes to metabolite concentrations through turnover-rate constraints while preserving the modularity of constraint-based approaches (Figure 1).

METEOR-K supports both global and targeted applications. Turnover-rate constraints can be imposed network-wide using physiologically relevant reference timescales or selectively on metabolite pools identified by modal analysis as being associated with slow dynamical modes. As part of the framework, we also developed turnover-aware sampling strategies to efficiently explore the resulting turnover-constrained feasible space.

We evaluate METEOR-K across network topologies and scales, from a reduced glycolysis pathway to near-genome-scale ovarian cancer metabolism, with complementary *E. coli* applications under anaerobic and aerobic conditions. Together, these applications show that METEOR-K provides a scalable route for screening steady-state metabolic states for turnover-based dynamic plausibility and prioritizing the constraints most relevant to local metabolic timescales. By restricting turnover-inconsistent states before downstream kinetic analysis, METEOR-K provides a more physiologically grounded starting point for model-guided exploration of metabolic engineering interventions.

## Results

### Turnover rates shape accessible local timescales

Using a reduced glycolysis model, we tested whether turnover rates computed from steady-state flux-concentration pairs can identify metabolite pools that limit attainable local dynamics. We quantify flux-to-concentration coupling by defining the turnover rate of metabolite *i, TR*_*i*_, as:

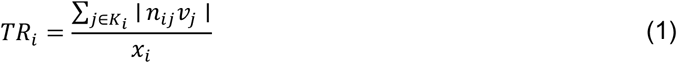

where *x*_*i*_ is the metabolite concentration, *v*_*j*_ is the flux through reaction *j, n*_*ij*_ is the stoichiometric coefficient of metabolite *i* in reaction *j*, and *K*_*i*_ is the set of reactions that consume metabolite *i* at the considered steady state. Thus, *TR*_*i*_ has units of inverse time and quantifies how rapidly a metabolite pool is renewed relative to its size. Equivalently, 1/*TR*_*i*_ is the characteristic time required to replace that pool at the given steady-state fluxes. At steady-state, total production and total consumption are equal in magnitude, so the turnover can be computed from either the producing or consuming flux set.

The link between turnover and local dynamics follows from the structure of the Jacobian: its entries scale with flux-to-concentration ratios, while reaction-rate sensitivities and network coupling determine how individual modes are realized (Supplementary Notes 1 and 2). Thus, turnover rates provide an indicator of the timescale range accessible to the linearized system. In particular, when the Jacobian is weakly coupled or diagonally dominant, turnover rates can track the eigenvalue scale and therefore the attainable local dynamic response. In strongly coupled networks, however, turnover rates influence the distribution of accessible local timescales rather than directly determining individual eigenvalues. These eigenvalues depend on how flux-to-concentration scaling combines with reaction-rate sensitivities and off-diagonal interactions in the Jacobian.

We illustrate the effects of the turnover rate using a reduced glycolysis model (Figure 2A). The network describes glucose uptake, conversion to glucose-6-phosphate (g6p), a lumped glycolytic step to phosphoenolpyruvate (pep), and subsequent conversion to pyruvate (pyr), which then branches toward lactate and acetyl-CoA (accoa). This branching structure creates a potential slow pool, because accoa can receive only a fraction of upstream carbon flux while remaining within its feasible concentration range. To test whether local dynamic behavior is shaped by the underlying flux-concentration state, we sampled 5,000 feasible steady states spanning the network’s solution space and built an ensemble of 100 kinetic models consistent with each sample using the ORACLE framework ^24^, retaining only dynamically stable models (Methods). We converted the eigenvalue with the largest real part of each stable kinetic model into a characteristic relaxation time and used the median characteristic time across models to summarize each steady state. Different regions of the steady-state solution space corresponded to markedly different characteristic times, producing a gradient across the PCA projection of sampled states (Figure 2B). The same trend was observed when using the fastest sampled characteristic time for each steady state (Figure S1), indicating that the effect does not depend on the aggregation metric.

**Figure 2.**
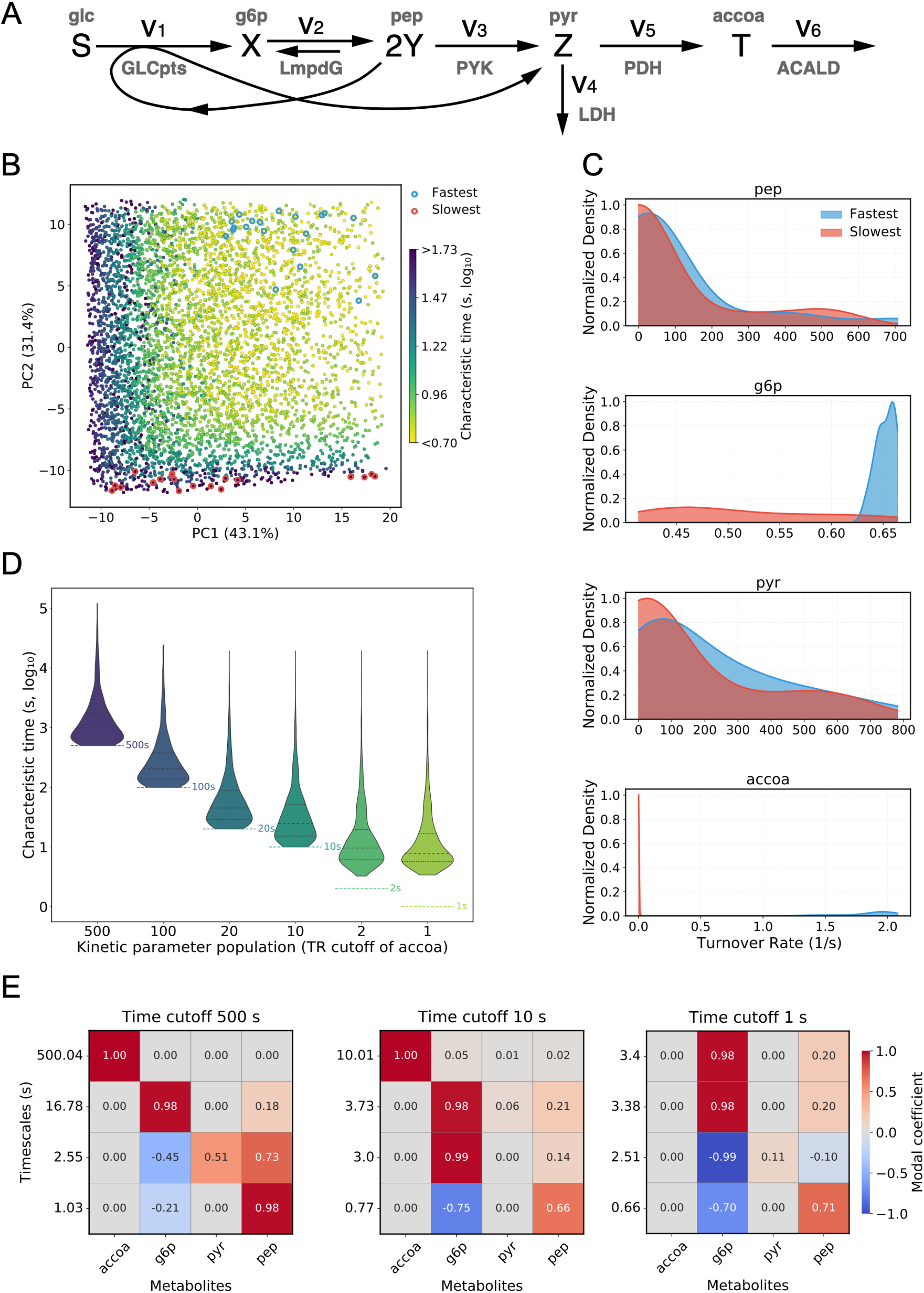
Turnover-rate constraints shift local timescales in a reduced glycolysis model. (A) Schematic of the reduced glycolysis pathway with metabolites and fluxes (V1–V6) connecting glc (S), g6p (X), pep (2Y), pyr (Z), and accoa (T). The pyruvate branch creates a potential low-turnover accoa pool. (B) Principal component analysis (PCA) of 5,000 feasible steady-state samples. Points are colored by the median characteristic time (log_10_) computed from 100 kinetic models parameterized around each steady state. Blue circles mark the 20 fastest steady states, and red circles mark the 20 slowest steady states by median characteristic time. (C) Turnover rate distributions (1/s) for pep, g6p, pyr, and accoa in the fastest and slowest steady-state groups from panel B. Each subplot shows normalized density for one metabolite; blue indicates the fastest states and red indicates the slowest states. Accoa shows the strongest separation between slow and fast states. (D) Characteristic time distributions for kinetic model populations generated while imposing different accoa turnover time cutoffs. The y-axis shows characteristic time in seconds (log_10_), and colored dashed horizontal lines indicate the corresponding accoa turnover time cutoff. (E) Modal analysis linking dominant timescales to metabolite pools. Heat maps show modal coefficients for the fastest kinetic models under representative accoa TR time cutoffs (500 s, 10 s, and 1 s). Reported values show the contribution of each metabolite to the mode associated with the indicated timescale.

To identify which metabolite pools were associated with these differences, we compared the turnover rate distributions of the 20 fastest (Figure 2C, blue) and 20 slowest (Figure 2C, red) steady states. Pep and pyr showed overlapping distributions, and g6p displayed only partial separation. In contrast, accoa showed a pronounced shift: slow steady states were consistently associated with very low accoa turnover, whereas fast states exhibited substantially higher values. Thus, in this simple network, the metabolite with the lowest turnover sets the slowest accessible local timescale. This correspondence is expected from the network structure, because accoa lies in a simple linear branch directly linked to the turnover of that pool, as shown analytically in Supplementary Note 2.

We next tested how increasing accoa turnover affected local dynamics. We constrained the turnover rates of all metabolites to a characteristic time of 2 s, except for accoa, which we systematically varied from 500 s to 1 s. For each case, we generated a representative steady state, sampled 1,000 kinetic models around it, and quantified the resulting characteristic times (Figure 2D). As accoa turnover increased, the system’s fastest attainable dynamics accelerated accordingly, until accoa no longer represented the slowest pool. At 2 s and 1 s of accoa’s turnover time, g6p became the new bottleneck. Modal analysis confirmed this bottleneck shift: the slowest mode was associated with accoa when its turnover remained slow and with g6p once accoa turnover was increased (Figure 2E). This analysis indicates that slow turnover of a specific metabolite pool can limit the fastest accessible local timescale in this reduced network.

Finally, when we pushed all metabolite turnover rates toward their highest feasible values, the system approached a theoretical upper bound on attainable local dynamics, jointly set by thermodynamic constraints, mass balance, and flux bounds (Supplementary Note 3). Together, these results show that steady-state flux-concentration pairs already encode constraints on attainable kinetic timescales. Turnover rates, therefore, provide an interpretable way to identify metabolite pools that make otherwise feasible steady states dynamically slow. We next formalize this principle as a mixed-integer optimization problem to identify turnover-constrained, dynamically plausible steady-state metabolic states.

### Formulation of turnover-constrained, dynamically plausible metabolic states

METEOR-K constrains steady-state metabolic states by combining mass-balance constraints, thermodynamic feasibility, and metabolite turnover-rate constraints into a single mixed-integer linear programming (MILP) optimization problem. The feasible steady-state flux space is first defined by mass-balance, flux bounds and any additional linear constraints from measured exchange rates or data-integration procedures:

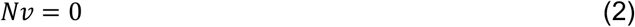

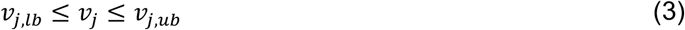

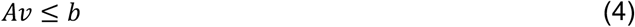

where *N* ∈ ℝ^*m*×*n*^ is the stoichiometric matrix, *v* ∈ ℝ^*n*^ the vector of reaction fluxes, and Eq. 4 stems from additional physiological or data-derived constraints.

Thermodynamic feasibility is imposed using the standard Thermodynamics Flux Analysis (TFA) ^11^ formulation, which couples reaction directionality to transformed Gibbs free energies and metabolite concentrations through mixed-integer constraints. For each metabolite *i*, METEOR-K uses log-transformed concentrations

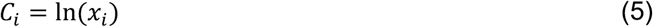

with bounds

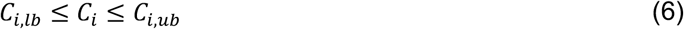

For each reaction *j*, the reaction Gibbs free energy is linked to the log-concentrations by

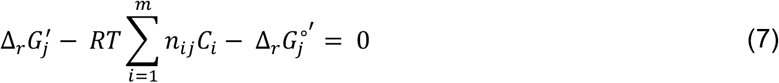

where *R* is the gas constant, *T* is the temperature, and *n*_*ij*_ is the stoichiometric coefficient of metabolite *i* in reaction *j*. Reaction directionality is then constrained by the sign of 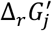, using the standard TFA binary-variable formulation described in the Methods. These thermodynamic constraints exclude flux states that violate reaction energetics, but they do not constrain how fast metabolite pools are replenished. Thus, two steady-state solutions can satisfy the same mass-balance and thermodynamic constraints while differing substantially in metabolite pool size relative to consuming flux. Large metabolite pools relative to their consuming fluxes imply slow turnover and potentially implausible metabolic timescales.

METEOR-K addresses this limitation by requiring selected metabolites to satisfy turnover-rate bounds. For metabolite *i*, we define the total consuming flux as:

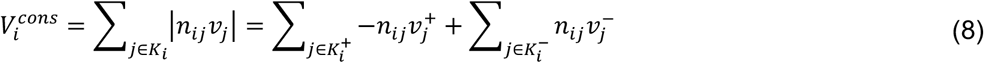

where *K*_*i*_ is the set of reactions that consume metabolite *i* at the considered steady state. The flux variables can be split into non-negative variables representing the forward 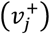 and reverse directionality 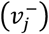. 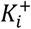 contains reactions that consume metabolite *i*, i.e.,*n*_*ij*_ < 0, whereas 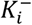 contains reactions for which reverse flux consumes metabolite *i*, i.e., *n*_*ij*_ > 0. This representation makes 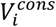 linear in the split-flux variables 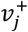 and 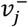. We impose metabolite-specific turnover intervals of the form

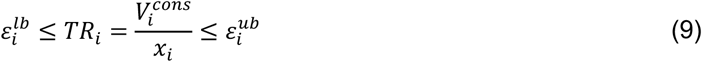

where 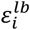 and 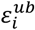 define lower and upper physiological turnover-rate bounds for metabolite *i*. Combining Eq. 5 and Eq. 9 yields:

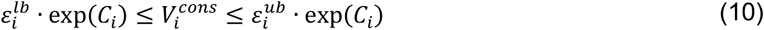

These constraints are nonlinear and, when combined with the binary variables used to enforce reaction directionality, direct implementation would produce a mixed-integer nonlinear programming problem. To preserve computational scalability and retain a MILP formulation, METEOR-K approximates the exponential term using either concentration-bin variables or a piecewise-linear approximation over the feasible log-concentration range. The full formulation, including thermodynamic constraints, turnover-constraint linearization, approximation-error analysis, and computational trade-offs, is described in the Methods.

### Turnover rate constraints restrict the steady-state space and accelerate metabolic response times in anaerobic *E. coli*

We next moved beyond the reduced network to test whether turnover-rate constraints restrict the admissible steady-state space, reduce flux-concentration uncertainty, and select states with faster local and nonlinear metabolic responses in a larger metabolic model. We tested this formulation on the published anaerobic *E. coli* metabolic model ^25^. The stoichiometric network contains 180 reactions and 141 metabolites, and the associated kinetic model includes 63 ODEs and 478 kinetic parameters (Methods). We constrained the model to the experimentally measured anaerobic growth rate and imposed lower bounds on metabolite turnover rates using the cell doubling time as a reference timescale. Because these bounds define minimum turnover rates, metabolites whose feasible turnover rates were already higher than the inverse doubling-time reference remained effectively unconstrained. For example, in this *E. coli* model, ATP turnover could be more than 200-fold faster than cell doubling.

We generated 100 alternative steady-state solutions for increasingly stringent lower bounds on TR, corresponding to timescales 5-, 10-, 50-, and 100-fold higher than the cell’s doubling time (Figure 3A). For each steady-state solution, we sampled 100 kinetic parameter sets and computed the corresponding characteristic times. The resulting steady states occupied distinct regions of the feasible space in the PCA projection (Figure 3A) and, on average, exhibited different linear dynamics (Figure 3B). As TR lower bounds became increasingly tighter, both the fastest attainable and median characteristic times decreased. However, the median characteristic time plateaued at approximately the 10-fold cutoff, whereas the fastest achievable characteristic time continued to decrease as the cutoff became stricter (Figure 3B). This pattern indicates that TR constraints shift the system’s accessible timescales, but beyond a threshold, the median timescale is influenced more by kinetic-parameter variability, thermodynamic constraints, and network topology than by the global TR cutoff.

**Figure 3.**
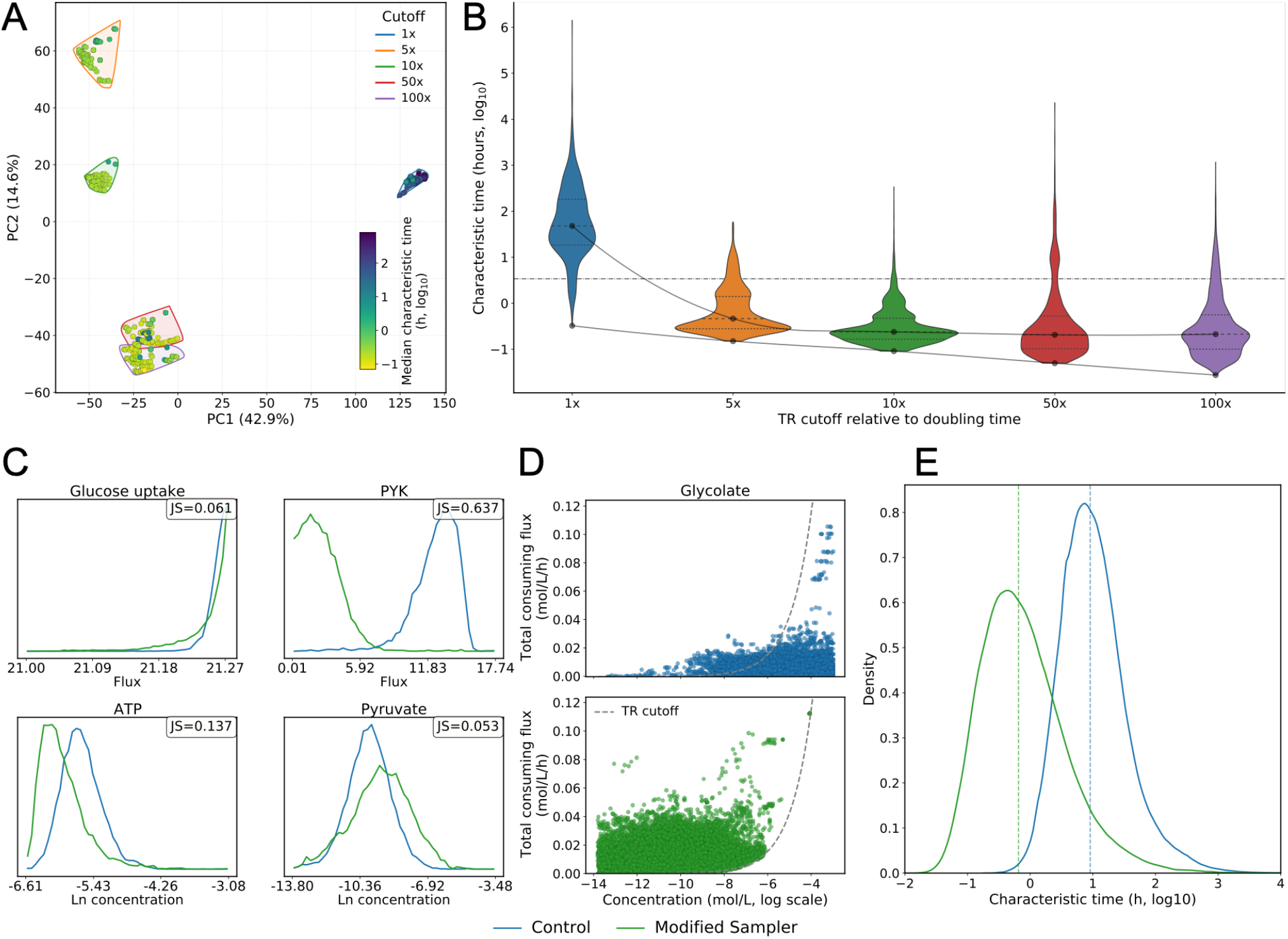
METEOR-K restricts steady-state flux-concentration space and accelerates local dynamics in an anaerobic *E. coli* model. (A) Principal component analysis of 100 alternative steady-state solutions generated with a global TR lower bound applied to all metabolites at 1-, 5-, 10-, 50-, and 100-fold faster than the cell doubling time. Points are colored by the median characteristic time (log_10_) computed from kinetic models parameterized around each steady state. (B) Violin plots of characteristic-time distributions for kinetic-model populations generated from steady states in each TR-cutoff group. Black dots mark medians, and dashed lines mark the 25th and 75th percentiles. (C) Representative marginal distributions from 20,000 steady-state samples generated with a TR-aware sampler (green) and a control sampler without TR constraints (blue). Jensen-Shannon divergence values quantify the distributional shift for each variable. (D) Flux-concentration plane for glycolate, showing total consuming flux versus sampled concentration. The dashed curve indicates the imposed TR cutoff boundary; blue points show control samples and green points show TR-aware samples. (E) Characteristic-time distributions for kinetic models generated from the TR-aware and control steady-state sample sets; dashed lines indicate medians.

These results show that TR lower bounds shift both the location of feasible steady states and the distribution of accessible linear timescales. To quantify how strongly TR constraints reshape the broader admissible flux-concentration space, we compared TR-constrained and unconstrained steady-state ensembles using turnover-aware sampling, applying a global TR lower bound corresponding to turnover times 10-fold shorter than the cell doubling time (Methods). TR constraints produced substantial changes in the sampled distributions: the average Jensen-Shannon (JS) divergence (computed with the natural logarithm) was 0.301, corresponding to approximately 43% of the theoretical maximum, and only 27% of variables had JS divergence ≤ 0.1, indicating substantial shifts in the sampled distributions. Representative examples are shown in Figure 3C.

To understand which parts of the network drove these distributional shifts, we compared extracellular exchange fluxes, intracellular fluxes, and metabolite concentrations between unconstrained and TR-constrained samples. Extracellular exchange fluxes were largely unchanged: glucose uptake and, more generally, secretion and uptake rates showed similar distributions across the two sampling regimes (Supplementary Table S1). This limited change likely reflects, in part, the model’s prior integration of experimentally measured uptake and secretion rates, although not all exchange fluxes were directly constrained by these data. In contrast, many intracellular pathways differed markedly. For example, fluxes in pyruvate metabolism had a median JS divergence of 0.258. PYK, which catalyzes the terminal pyruvate-producing step of glycolysis, showed markedly lower sampled fluxes under TR constraints (Figure 3C). This decrease was accompanied by higher DHAPT flux, while total pyruvate-consuming flux remained nearly unchanged. Pyruvate concentrations increased, resulting in slightly lower pyruvate turnover rates that nevertheless remained above the imposed lower bound. More broadly, most cytosolic metabolites showed lower sampled concentrations under TR constraints, consistent with the exclusion of states containing large metabolite pools relative to their consuming fluxes.

TR constraints also reduced uncertainty in intracellular metabolite concentrations by excluding high-pool/low-consuming-flux regions that remain feasible under mass-balance and thermodynamic constraints alone. To make this exclusion explicit, we plotted, for each metabolite, its total consuming flux against its sampled concentration. An imposed TR lower bound defines a clear exclusion region in the flux-concentration plane, corresponding to metabolite pools too large to be renewed by the available consuming flux. Glycolate illustrates this behavior: TR-constrained samples occupied a distinct region from unconstrained samples (Figure 3D), and the same qualitative pattern was observed for other metabolites. To quantify the associated reduction in marginal uncertainty across the network, we used thermodynamic variability analysis to compute the feasible ranges of all fluxes and concentrations with and without TR constraints and expressed the effect as the fractional reduction in the feasible range. For the 10-fold TR lower bound cutoff, concentration ranges showed measurable reductions, whereas flux-range changes were negligible (Supplementary note 6). This asymmetry is informative: in this model, fluxes were already strongly constrained by stoichiometry, thermodynamics, and measured exchange rates, whereas metabolite concentrations retained broader feasible ranges. Adding TR constraints, therefore, primarily refined the admissible concentration space. For example, the GTP concentration range was reduced by 69%.

When we sampled 100 kinetic parameter sets for each of the two steady-state ensembles, the resulting kinetic models differed markedly in their dynamical timescales. Models generated from TR-constrained steady states exhibited an approximately one order of magnitude shorter median characteristic time, i.e., faster local dynamics (Figure 3E).

To ensure that these distributional changes arise from the TR constraints rather than from artifacts of the modified sampling procedure, we compared the TR-aware sampler against a baseline approach that uses a standard sampler with accept-reject filtering based on the same TR constraints. The two approaches produced nearly identical distributions for all variables, with an average JS divergence of 0.0308 and more than 95% of variables showing JS divergence ≤ 0.1. This agreement suggests that the modified sampler does not introduce additional sampling bias beyond enforcing feasibility under TR constraints.

We next tested whether constraining metabolite turnover rates also affects system behavior in a nonlinear setting using batch bioreactor simulations. We simulated biomass accumulation by coupling each kinetic model to an exponential biomass-growth equation and parameterized the initial batch conditions using experimental data ^26^. To generate the kinetic models used in these simulations, we sampled steady states under TR constraints corresponding to turnover times equal to the doubling time and 5- and 10-fold shorter than the doubling-time reference. To ensure favorable initial conditions for the reactor simulations, we enforced high steady-state concentration ranges for biomass precursors identically across all TR cutoffs, so that differences between cutoffs reflect the turnover constraint rather than the precursor enrichment (Supplementary Note 4).

We integrated each batch bioreactor simulation for 15 hours; in the reference experimental setting, cultures reached the stationary phase at approximately 10.5 hours. As the primary readout, we report the average growth rate, which is directly comparable to the experimentally measured rate, and treat the biomass concentration at 10.5 hours as a complementary check. Outcomes varied substantially across simulations, attributable to both the steady-state sample used to initialize the inoculum and the sampled kinetic parameter values governing carbon and energy flow through the network. Despite this variability, increasing the TR cutoff consistently increased the emergent average growth rate (Figure 4A). The experimentally measured anaerobic growth rate is shown as a physiological reference but was not used to fit or select the TR cutoff. Rather, the result shows that steady states constrained to faster metabolite turnover yield kinetic models with faster growth dynamics in nonlinear bioreactor simulations. Biomass titer followed the same ordering (Figure 4B), but with greater scatter because titer also depends on the initial biomass associated with each steady-state sample. This trend persisted after stratifying simulations by their initial maximum eigenvalue, indicating that the TR lower bound, rather than the initial characteristic time alone, explains the increase in biomass titer (Figure 4C).

**Figure 4.**
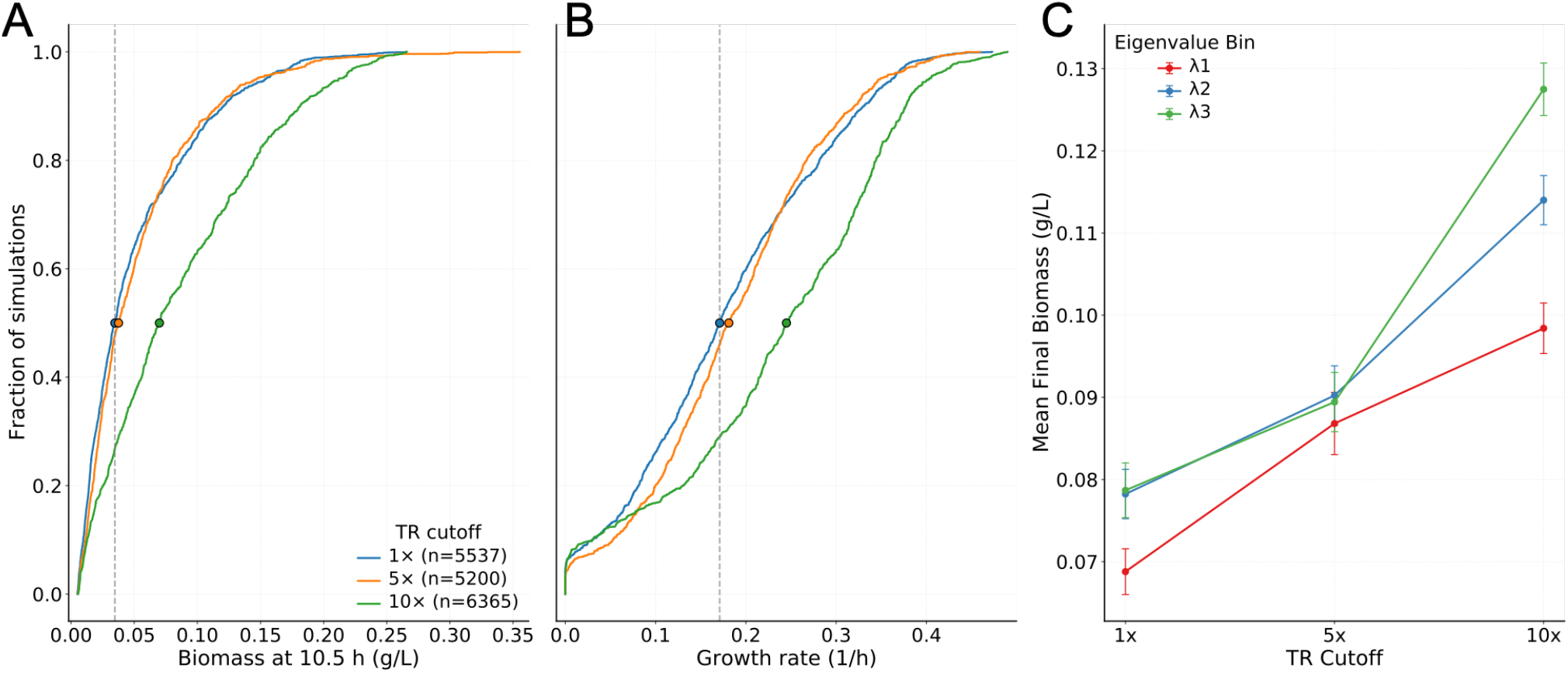
TR constraints accelerate nonlinear growth dynamics in anaerobic *E. coli* bioreactor simulations. (A-B) Empirical cumulative distributions of average growth rate in bioreactor simulations (A) and biomass concentration at 10.5 h (B) for kinetic models initialized from steady states generated with global TR cutoffs of 1-, 5-, and 10-fold shorter than doubling time. Circles mark medians, and the number of simulated models is given in the legend. The dashed vertical line in (A) indicates the experimentally observed anaerobic growth rate as a physiological reference. (C) Average final biomass grouped by TR cutoff and initial eigenvalue bin. Error bars indicate the standard deviation within each group.

Together, these results show that turnover constraints that accelerate local relaxation timescales also shift emergent nonlinear growth behavior, providing a biophysically interpretable connection between steady-state flux-concentration structure and dynamic phenotype.

### Targeted turnover-rate enforcement scales to a multi-omics-constrained cancer metabolic model

We next asked whether METEOR-K can be applied in a large-scale metabolic network whose feasible space is already tightly constrained by thermodynamics and multi-omics data. This setting is important because a single stringent TR lower bound applied uniformly across all metabolite pools can conflict with existing data-derived and thermodynamic constraints. We therefore used a large-scale ovarian cancer metabolic model developed in our previous work^27^ as an example of such a network and tested the targeted mode of METEOR-K, in which turnover constraints are imposed only on metabolite pools associated with slow relaxation modes. The ovarian cancer model had already been constrained by thermodynamic feasibility and by multi-omics data integration, including metabolomics, exchange-flux measurements, and transcriptomics (Methods). These constraints defined a reduced feasible space of fluxes and concentrations consistent with the available biochemical and experimental evidence. The model has broad network coverage, with 2137 intracellular reactions and 974 metabolites spanning 8 compartments, and the accompanying kinetic model comprises 716 ODEs and 9594 kinetic parameters.

Within this already restricted feasible space, enforcing a sufficiently stringent TR lower bound uniformly across all metabolite pools would overconstrain the current data-integrated formulation. Restoring feasibility would require changing the model assumptions, for example by lowering concentration bounds for metabolites with slow-consuming fluxes or by relaxing thermodynamic constraints that restrict the feasible concentration space. Because these bounds and constraints encode the integrated data and thermodynamic prior information, we retained them unchanged and instead applied targeted, modal-analysis-guided turnover enforcement. This strategy applies turnover constraints where they are most dynamically informative, rather than uniformly across the network (Figure 1, Methods).

The targeted workflow is iterative (Figure 1). We first generate an initial set of steady-state samples and parameterize kinetic models around each sample, allowing characteristic times and modal contributions to be computed. We then impose TR lower bounds on the metabolite pools most consistently associated with slow modes. Because this identification is statistical and depends on the current ensemble, the procedure is repeated for a few iterations until the set of slow modes stabilizes or the median characteristic time improves only marginally (Methods).

Applying this iterative procedure to the cancer model, modal analysis identified 705 metabolite pools that repeatedly contributed to slow modes. We imposed a TR lower bound corresponding to a turnover of 15-fold shorter than the cell doubling time. This progressively shortened the median characteristic time of the kinetic-model ensembles to nearly an order of magnitude below the unconstrained baseline (Figure 5A). The TR lower bound is a tunable parameter, and we recommend a stricter cutoff for large networks because the probability of sampling dynamically favorable steady states falls as network size grows. Raising the lower bound biases the feasible region toward states with faster local turnover and increases the chance that randomly parameterized kinetic models initialized at these states exhibit acceptable timescales.

**Figure 5.**
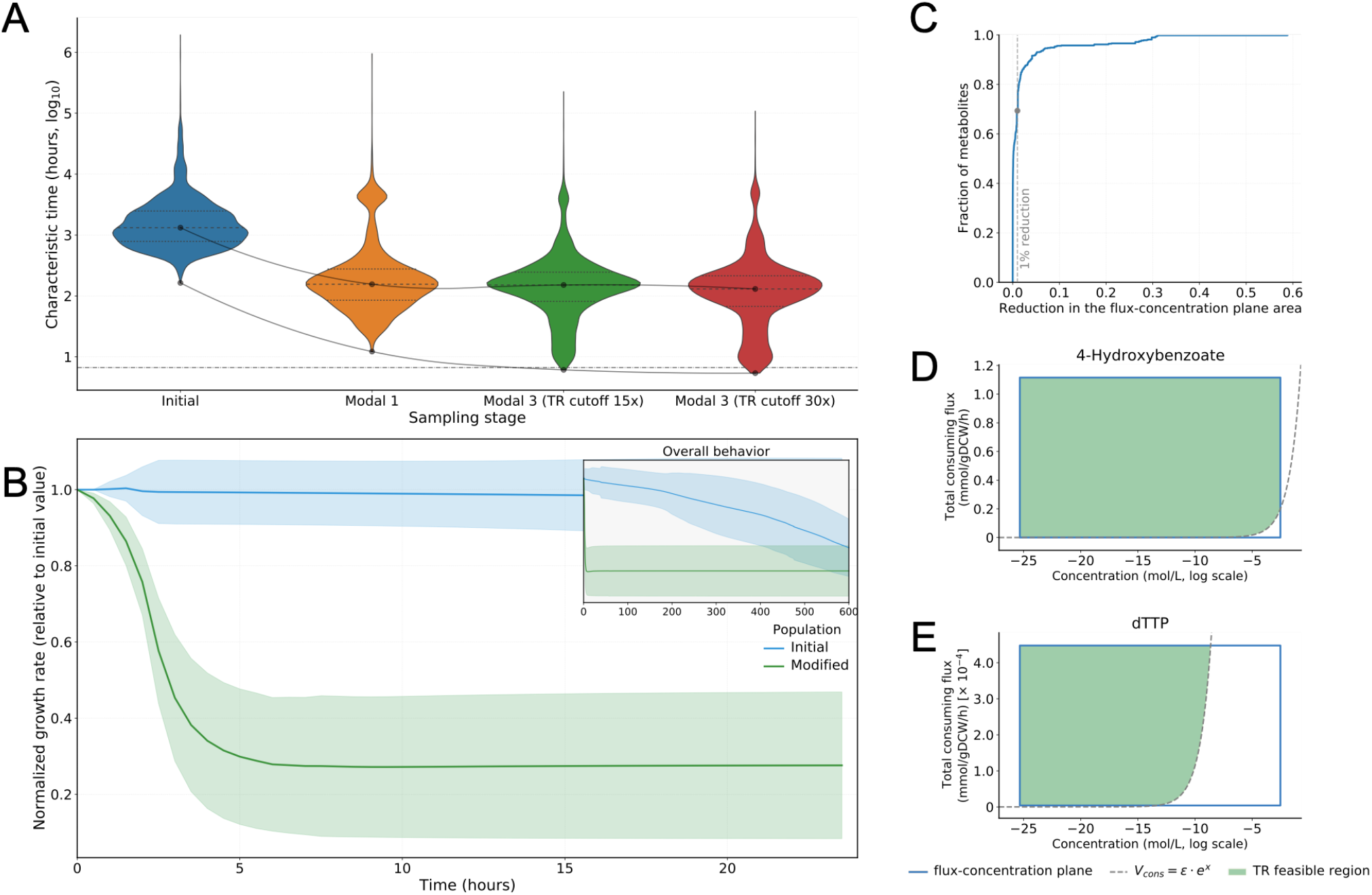
Targeted TR enforcement improves local dynamics and accelerates responses in nonlinear ODE simulations in the ovarian cancer model. (A) Violin plots show characteristic time distributions for kinetic model populations generated from steady states at different iterations of the modal analysis workflow. Black dots mark the medians, and dashed lines mark the 25th and 75th percentiles. The green and red distributions use the same targeted metabolite set but different TR lower bounds, corresponding to turnover times 15- and 30-fold shorter than the cell doubling time, respectively. (B) Dynamic drug-administration simulations in which thymidylate synthase activity was reduced, and growth-rate trajectories were followed from initial and TR-modified kinetic model populations. Lines show the mean normalized growth rate relative to the initial value, and shaded regions show one standard deviation. The inset shows the same trajectories over the full 600 h simulation horizon. (C) Empirical cumulative distribution of fractional reductions in flux-concentration plane area across constrained metabolites after applying targeted TR constraints. (D-E) Flux-concentration planes for representative metabolites. The dashed curve marks the TR boundary, with the green shading indicating the TR-feasible region retained after imposing the constraint.

Most of this improvement was achieved within the first two iterations: beyond the second, the median characteristic time no longer improved appreciably, although the fastest sampled characteristic times continued to decrease. This plateau indicated that the 15-fold cutoff, rather than the number of iterations, had become the limiting factor. Thus, additional iterations at the same cutoff would yield only diminishing improvements in the median characteristic time. We therefore raised the TR lower bound for the 705 identified metabolites from 15- to 30-fold, which shifted the ensemble toward faster attainable dynamics (Figure 5A). Applying a pruning criterion that required the characteristic time to be at least 3 times faster than the cell doubling time, 169 kinetic models (from 14 of 100 steady-state samples) were eligible at the 15-fold TR cutoff and 367 models (from 23 of 100 samples) at the 30-fold TR cutoff. Thus, once additional iterations at the same cutoff produced diminishing returns, increasing the TR lower bound improved the yield of dynamically acceptable kinetic models.

### Targeted turnover-rate enforcement enriches cancer-model ensembles for nonlinear enzyme-perturbation recovery and reduces the feasible flux-concentration space

TR constraints do not create dynamic behavior absent from the original feasible space. Rather, they exclude flux-concentration states with necessarily slow pool renewal and bias sampling toward regions more likely to yield physiologically relevant dynamics after kinetic parameterization. The dynamics ultimately realized still depend on kinetic parameters, elasticities, and network topology, which TR does not determine. In principle, unconstrained sampling can also generate steady states with favorable dynamic properties; in practice, this becomes increasingly unlikely in large networks, where the feasible space is high-dimensional and most unconstrained samples fall in regions associated with slow local dynamics, as illustrated by the initial distribution in Figure 5A. We therefore asked whether kinetic models parameterized from the unconstrained sampling distribution provide suitable starting points for nonlinear simulations.

To evaluate this, we reduced thymidylate synthase (TMDS) activity exponentially by 10-fold over an 8-hour window and then maintained it at the reduced level for the remainder of the simulation (Methods). We integrated each kinetic model for 600 simulation hours, allowing both the growth-rate reaction and intracellular metabolism to adapt to a new operating steady state. We compared the 169 TR-constrained models obtained after the third iteration that also satisfied the characteristic-time pruning criterion with 150 models drawn from the unconstrained ensemble, selected to span the original characteristic-time distribution. Because no unconstrained models satisfied the same pruning criterion, this comparison tests unconstrained models across their available timescale range rather than as a matched fast-timescale control.

We monitored the growth-rate reaction and recorded the time required to reach a new steady value. The TR-constrained models typically stabilized within a few hours (Figure 5B). In contrast, none of the unconstrained models reached a stable growth rate within 600 simulation hours, including those drawn from the fastest available part of the unconstrained distribution (Figure 5B). Thus, targeted TR enforcement enriched the ensemble for recovery-capable models that were not accessed by unconstrained sampling.

To assess convergence of the whole intracellular network, we also applied an equilibrium-detection algorithm to determine when all intracellular concentrations and fluxes reached a new steady state (Methods). Using this criterion, 60% of the TR-constrained models achieved a complete intracellular steady state within 600 (simulation) hours, whereas none of the unconstrained models did, consistent with the slow growth-rate trajectories shown in Figure 5B. Targeted TR enforcement therefore improved both output-level recovery of the growth rate and, for a subset of models, whole-network equilibration.

We next quantified how much the TR constraints reduced the feasible solution space of the human cancer model. Starting from the original model with baseline flux and concentration bounds, we imposed TR constraints on all metabolites identified up to the third iteration of the modal-analysis workflow, using the 15-fold lower-bound cutoff. We then performed variability analysis to estimate the feasible ranges of all concentrations and fluxes. As an aggregate view of this targeted 15-fold enforcement, we also estimated the contraction of the joint feasible flux-concentration space using a bounding-box approximation based on the flux and concentration ranges obtained from variability analysis (Supplementary Note 6).

To assess the local effect of each TR constraint, we calculated the area of the total consuming flux-concentration plane for each constrained metabolite before and after TR enforcement. Specifically, this plane was defined by the metabolite’s feasible concentration range and by the feasible range of total consuming flux at each concentration value. We estimated the fractional reduction in this plane by comparing the feasible area retained after imposing the TR boundary with the original area defined by the baseline variability ranges. This metric summarizes how strongly the TR constraint restricts the joint flux-concentration combinations available to each metabolite.

Across the constrained metabolites, most individual reductions were small (Figure 5C). Approximately 70% of metabolites showed less than a 1% reduction in their flux-concentration plane, 25% showed reductions between 1% and 10%, and 5% showed reductions greater than 10%. Thus, for most metabolites, the imposed TR constraint removed only a small fraction of the local flux-concentration plane. Yet, their cumulative effect across the solution space was reflected in shifted characteristic-time distributions and improved nonlinear recovery.

Figures 5D and 5E show representative examples of how TR constraints reduce flux-concentration planes for individual metabolites. The feasible concentration ranges for 4-hydroxybenzoate and dTTP were similar, but their feasible consuming flux ranges differed by approximately four orders of magnitude. For 4-hydroxybenzoate, the consuming-flux range was high enough that the TR constraint was nearly redundant and only combinations of very low consuming flux and high concentration violated the TR lower bound (Figure 5D). In contrast, dTTP had a much lower consuming flux range, so the TR constraint substantially reduced its feasible concentration range (Figure 5E). This behavior indicates that metabolites in low-activity regions of the network cannot simultaneously maintain large pool sizes and satisfy fast turnover requirements.

Together, these results demonstrate that targeted turnover-rate enforcement can preserve a tightly data-constrained cancer metabolic phenotype while enriching kinetic-model ensembles for physiologically compatible responses to enzyme perturbation. The ovarian cancer application therefore establishes the scalability and generality of METEOR-K in a complex metabolic network and provides turnover-consistent reference states for subsequent model-guided analysis of metabolic interventions.

### Data-informed turnover constraints restrict feasible states and reshape kinetic ensembles

The preceding applications used turnover cutoffs defined relative to cell doubling time. We next assessed how literature-derived, metabolite-specific turnover intervals restrict feasible metabolic states and influence the kinetic ensembles constructed around them. We derived turnover-time intervals for 18 central-carbon metabolites from experimental flux and concentration data reported in four *E. coli* studies (Supplementary Table S2) ^28–31^. These intervals collectively spanned 0.017 to 41.1 s, with the largest within-pathway fold variation observed among TCA-cycle metabolites (Figure 6A).

**Figure 6.**
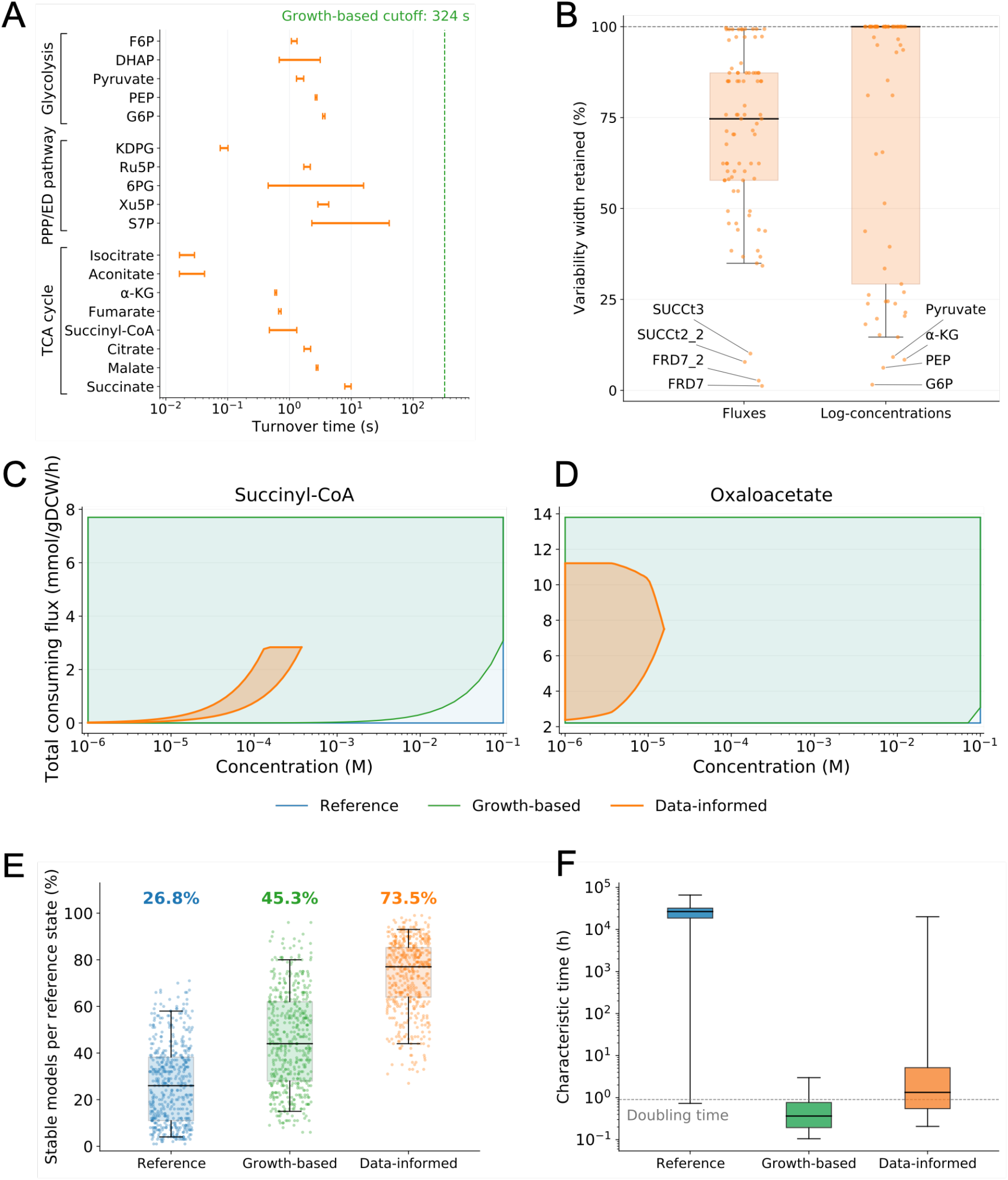
Data-informed turnover constraints restrict feasible metabolic states and alter kinetic ensembles. (A) Literature-derived turnover-time intervals for 18 metabolites, assembled from four studies. The green dashed line marks the growth-based cutoff. (B) Reaction-flux and log-concentration variability widths in the data-informed case, expressed as percentages of the corresponding reference-case widths. Each point represents one variable reaction flux or log-concentration. Counts denote variables with range reductions greater than 1%. (C, D) Feasible total-consuming-flux– concentration projections for succinyl-CoA and oxaloacetate. (E) Fractions of locally stable kinetic models for each reference state and case. Dots denote reference states, and labels denote the overall stable fraction for each case. (F) Characteristic relaxation times for locally stable kinetic models. The dashed line marks the cell doubling time. In panels E and F, boxes show medians and interquartile ranges, and whiskers span the 5th-95th percentiles.

We applied these turnover constraints to an aerobic *E. coli* core model comprising 89 reactions and 69 metabolites spanning the central carbon metabolism ^32^. Using the same base model, we considered three cases that differed only in the turnover constraints imposed. The reference case included no turnover constraints. The growth-based case, representing the absence of metabolite-specific turnover information, imposed a common TR lower bound network-wide, requiring turnover times no longer than one-tenth of the cell doubling time (Figure 6A). In the data-informed case, the turnover-time intervals were converted to metabolite-specific lower and upper TR bounds for the 18 metabolites with literature-derived intervals, while the remaining metabolites were left unconstrained. All other steady-state constraints were identical across the three cases. The feasible steady states sampled from each case were subsequently used as reference states for constructing the corresponding kinetic-model ensembles.

Data-informed turnover bounds restricted flux and concentration ranges throughout the network (Figure 6B). Among 86 reactions with variable reference flux ranges, 75 showed range reductions greater than 1%, with a median reduction of 25.3% across all 86 reactions. Log-concentration ranges narrowed by more than 1% for 32 of 65 variable pools, including 14 pools without direct turnover bounds. In contrast, the growth-based cutoff left the corresponding flux and concentration ranges essentially unchanged. The largest log-concentration-range reductions occurred for glucose 6-phosphate, phosphoenolpyruvate, α-ketoglutarate, and pyruvate, all of which had narrow imposed turnover-time intervals. Succinate-fumarate interconversion and succinate transport reactions showed the greatest flux-range reductions. Removing individual turnover constraints indicated that these four flux-reductions were driven primarily by the succinate bounds, with the fumarate bounds providing additional restriction.

Flux-concentration projections showed that turnover constraints restricted feasible states not only directly but also indirectly through network coupling. For succinyl-CoA, which had directly imposed data-informed turnover-rate bounds, the approximate feasible area of its total-consuming-flux–concentration projection was 96.8% smaller than in the reference case, compared with a 3.5% reduction in the growth-based case (Figure 6C). Oxaloacetate had no direct turnover bounds, yet its projected feasible area decreased by 85.4% in the data-informed case and by only 0.1% in the growth-based case (Figure 6D). The data-informed constraints reduced its maximum feasible concentration and narrowed the feasible range of its total consuming flux. This indirect restriction reflects propagation through network coupling, as oxaloacetate is linked through shared reactions to the turnover-constrained metabolites malate, citrate, and phosphoenolpyruvate.

To assess how the different turnover constraints affected the resulting kinetic ensembles, we sampled 500 distinct reference steady states for each case and parameterized 100 kinetic models around each state, yielding 50,000 models per case. The fractions of locally stable models were 26.8% for the reference ensemble, 45.3% for the growth-based ensemble, and 73.5% for the data-informed ensemble (Figure 6E). Among stable models, the median characteristic relaxation times were approximately 2.67 × 10^4^ h, 21.9 min, and 79.9 min for the reference, growth-based, and data-informed ensembles, respectively (Figure 6F). Thus, both turnover-constrained cases increased the fraction of stable models and shortened the median relaxation time relative to the reference case. The data-informed case yielded the highest stable fraction, whereas the growth-based case yielded the shortest median relaxation time.

We also considered a fourth, combined case in which the data-informed bounds were retained for the 18 metabolites and the growth-based TR lower bound was applied to the remaining metabolites represented in the kinetic model. Relative to the data-informed case, these additional constraints did not further narrow the reaction-flux or log-concentration ranges computed by variability analysis. Using the same steady-state-sampling and kinetic-parameterization procedure, 69.5% of the kinetic models were locally stable (Figure S5). Among stable models, the median characteristic relaxation time was 43.5 min, between the 79.9 min observed for the data-informed case and the 21.9 min observed for the growth-based case. The combined case contained fewer extremely slow models, shifting its characteristic-relaxation-time distribution toward that of the growth-based case.

Together, these results show that literature-derived turnover intervals can restrict feasible states beyond directly constrained pools and alter both the fraction of stable models and the relaxation-time distribution of the resulting kinetic ensembles. In this model, the growth-based constraint increased the stable fraction and shortened relaxation times without changing marginal reaction-flux and log-concentration ranges, whereas data-informed bounds also narrowed these ranges throughout the network. Extending the growth-based lower bound to the remaining metabolites reduced the prevalence of extremely slow models without further narrowing these marginal ranges.

## Discussion

METEOR-K addresses a missing link in steady-state constraint-based modeling: these models do not generally constrain whether flux magnitudes are compatible with the corresponding metabolite pool sizes, so admissible solutions can imply implausible pool-renewal timescales. By imposing turnover-rate constraints through a mixed-integer linear formulation, METEOR-K couples steady-state fluxes to metabolite concentrations and excludes flux-concentration states with pool-renewal timescales outside the selected physiological or experimental bounds. Across the systems studied here, these constraints restricted feasible flux-concentration states and shifted downstream kinetic models toward faster local dynamics and physiologically compatible nonlinear responses. METEOR-K thus provides a biologically interpretable addition to mass balance, thermodynamics, concentration bounds, and other data-derived constraints. Its turnover-rate constraints can be incorporated into constraint-based modeling workflows implemented with platforms such as the COBRA Toolbox ^33^.

Turnover constraints provide a physiological screen for dynamic plausibility by constraining flux-to-pool-size ratios that scale Jacobian entries. Low turnover indicates slow pool renewal, whereas high turnover permits, but does not guarantee, fast relaxation. A reversible reaction near equilibrium can nevertheless carry a small net flux while remaining highly sensitive to concentration perturbations, generating a fast local mode (Supplementary Note 2). Such a mode does not by itself imply fast whole-network relaxation. Other reactions must balance the metabolite pool: if they provide sufficient total throughput, the turnover bound is satisfied; if they operate with low flux-to-pool-size ratios, they may determine the dominant relaxation time. Turnover filtering thus evaluates pool renewal at the network level. Near-equilibrium sensitivity affects the relationship between turnover and dynamic timescales only when the resulting fast local mode is essential to the relevant dynamics and the balancing reactions neither raise total turnover above the cutoff nor introduce a slower dominant mode (Supplementary Note 2).

Consistent with this interpretation, progressively stricter TR lower bounds in the anaerobic *E. coli* model produced diminishing reductions in the median characteristic relaxation time. This result shows that turnover bounds shift the accessible timescale distribution, whereas kinetic parameters and network structure determine the dynamics realized within it. Turnover constraints can therefore enrich the reference steady-state ensemble for faster pool renewal and local dynamics, but kinetic parameterization and dynamic analysis remain necessary to determine the behavior actually realized. Because the extent of this enrichment depends on the selected cutoff, the constraints should be introduced progressively, beginning with an unconstrained baseline or a permissive cutoff and assessing how the results change as the cutoff increases. Where available, physiological reference timescales or experimental turnover intervals can provide a physiological or empirical basis for selecting the bounds and maintaining consistency with the modeled physiology.

How turnover constraints are applied in practice depends on the restrictions already imposed on the feasible space by data integration and thermodynamics. Under these conditions, a uniform global TR lower bound can render the formulation infeasible when no state simultaneously satisfies the data-derived, thermodynamic, and turnover constraints. Rather than relaxing the data-derived or thermodynamic priors, METEOR-K can enforce turnover constraints selectively on metabolite pools that modal analysis links to slow modes. In the cancer model, targeted enforcement preserved the existing data-derived and thermodynamic priors while yielding recovery-capable kinetic models that were not accessed by unconstrained sampling. METEOR-K therefore offers two complementary modes: global enforcement to filter turnover-inconsistent flux-concentration relationships and targeted enforcement to constrain the metabolite pools most relevant to slow local dynamics.

The aerobic *E. coli* application demonstrates how literature-derived turnover information can be incorporated directly into METEOR-K. Experimental turnover intervals can be obtained from isotope-labeling time courses ^34–36^, paired flux measurements and metabolite pool sizes ^29^, or published quantitative estimates of metabolite turnover times ^23,37,38^, and imposed as metabolite-specific bounds that reflect measurement uncertainty. Using intervals for 18 metabolites that encompassed reported uncertainty and variation across experimental conditions, METEOR-K narrowed reaction-flux and log-concentration ranges across the network, increased the fraction of locally stable models and shifted their relaxation-time distribution. These effects extended to metabolites without direct turnover bounds, showing how sparse turnover information can propagate through network coupling without imposing the underlying measurements as separate flux or concentration constraints.

METEOR-K complements enzyme-constrained modeling approaches such as GECKO ^16,39^ and ETFL ^15,40^, which use enzyme abundances, catalytic turnover numbers, and protein allocation constraints to limit fluxes. Whereas enzyme-constrained models restrict the catalytic capacity available to sustain reaction fluxes, METEOR-K links those fluxes to intracellular metabolite pool sizes through turnover constraints. Combining the two approaches could therefore restrict the feasible space to states that satisfy both enzyme-capacity and metabolite-pool-renewal requirements and identify inconsistencies between available catalytic capacity and experimentally supported turnover. Because METEOR-K is formulated as a MILP, it can, in principle, be integrated with enzyme-constrained frameworks that include or can be extended to include metabolite concentrations ^15,40,41^.

The reliability of kinetic-model-guided metabolic engineering depends not only on kinetic parameterization but also on the selection of reference steady states. Ensemble kinetic modeling workflows^42–45^ use sampled steady-state fluxes and metabolite concentrations for rate-law parameterization, stability analysis, and dynamic simulation. Rational-design frameworks such as NOMAD then use these ensembles to prioritize enzyme interventions.^46,47^ The value of this approach for strain design has been demonstrated experimentally, including improved xylose uptake following HXK2 deletion in *S. cerevisiae*^48^ and increased *p*-coumaric acid production in eight of ten engineered yeast strains^47^. These applications require sufficient network coverage and mechanistic detail as well as reference states consistent with the modeled physiology.

Hameri et al. showed that alternative flux-concentration states consistent with the same experimental physiology can yield substantially different metabolic-control estimates and engineering recommendations.^49^ While previous efforts have addressed parameter identifiability, observability, prediction uncertainty, and ensemble selection,^50–54^ METEOR-K addresses a distinct upstream source of uncertainty arising from reference-state selection by restricting the admissible flux-concentration space before kinetic parameterization, thereby providing turnover-consistent starting points for downstream intervention analysis. Whether turnover filtering also changes engineering-target rankings or improves the predictive accuracy of interventions remains to be established through further experimental validation.

Together, these results establish metabolite turnover as a tractable link between steady-state feasibility and physiologically grounded screening of metabolic timescales. By adding flux-to-pool-size constraints to standard steady-state formulations, METEOR-K supports constraint-based analysis and provides a scalable route to turnover-informed reference-state selection for kinetic modeling, strengthening the basis for rational metabolic engineering.

## Methods

### Implementation of turnover rate (TR) constraints

The turnover rate constraint defined in Eq. 10 was implemented in the TFA variable space, as described in the Results section. Because TFA uses log-concentration variables *C*_*i*_, and to keep the optimization formulation as an MILP, we represented the TR constraints in an MILP-compatible form. We imposed lower bounds in the growth-based and targeted applications and both lower and upper bounds in the experimental data-integration case study. The same linearization strategies can also be adapted to equality constraints. For simplicity, we present both discretization strategies below using the lower-bound constraint.

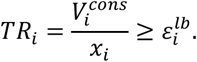

METEOR-K implements two MILP-compatible alternatives for this transformation: a concentration-bin-based approximation and a piecewise-linear approximation of the exponential concentration mapping.

#### Piecewise discretization based on concentration binning

For a metabolite *x*_*i*_ with a concentration range [*lb, ub*] we partition this range into N bins indexed by m = 1, …, N, with bounds [*lb*_*m*_, *ub*_*m*_]. Assume we impose a lower bound on the turnover rate of the metabolite 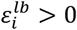. We also introduce binary variables *B*_*m*_ ∈ {0, 1} indicating whether bin *m* is active, auxiliary concentrations *AC*_*m*_, and log concentrations *LC*_*m*_. The nonlinear TR constraint from Eq. 10 can then be transformed to

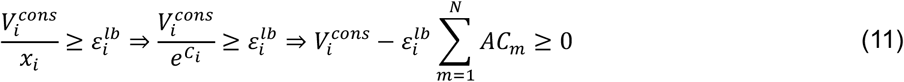

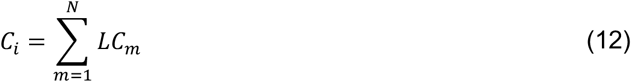

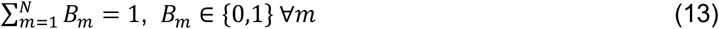

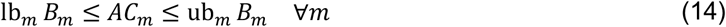

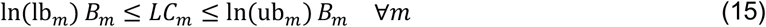

With this formulation, only one binary variable can be active at a time, constraining the values of the auxiliary log concentration variable *LC*_*m*_ and the approximation variable *AC*_*m*_. It is important to note that the active auxiliary variable *AC*_*m*_ does not necessarily equal the value implied by *C*_*i*_, because the bin constraints restrict only the admissible range within each interval and do not enforce an exact functional relationship between *AC*_*m*_ and *C*_*i*_. Therefore, the approximation error of this linearization depends on bin width and bin placement.

#### Piecewise-linear binning using SOS2-type constraints

We approximate the exponential mapping between log-concentration (*C*_*i*_) and concentration (*x*_*i*_) with a piecewise-linear construction enforced by SOS2-type constraints. We introduce breakpoints in log space *b*_1_ < *b*_2_ < ⋯ < *b*_*N*_ and define the corresponding function values 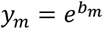. We also create weights *w*_*m*_ ∈ [0,1] such that {*w*_1_, …, *w*_*N*_} forms an SOS2 set, meaning that at most two adjacent *w*_*m*_ can be nonzero. Using this representation, the TR lower-bound constraint is imposed as

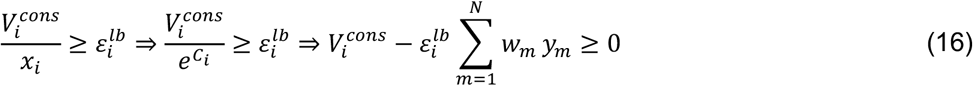

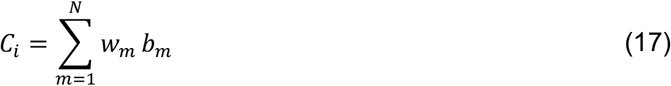

The SOS2 restriction was implemented by introducing binary variables *z*_*m*_ ∈ {0,1} for the *N* − 1 intervals between adjacent breakpoints. Exactly one interval can be selected, and each weight *w*_*m*_ is allowed to be nonzero only if one of its adjacent intervals is selected:

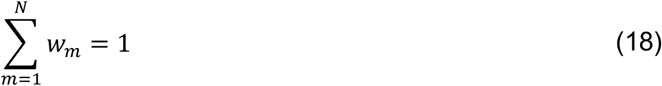

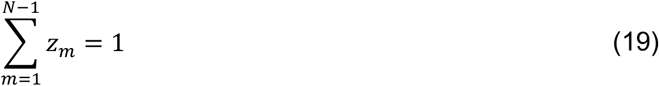

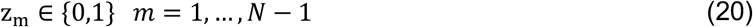

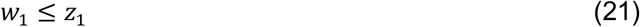

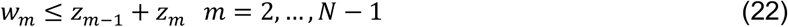

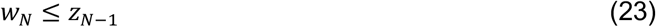

Because the exponential function is convex, its piecewise linear interpolation overestimates the true function. As a result, for lower-bound TR constraints, the approximated concentration is conservative with respect to the imposed constraint. The approximation error decreases as the number of breakpoints increases.

#### Approximation error in both techniques

The approximation error of the exponential mapping depends on the number and spacing of bins and breakpoints, the feasible concentration range, and the tolerated error relative to the imposed TR constraint. We quantify this error and provide guidance for selecting the number of bins, *N*, for both approaches in Supplementary Note 5.

#### Unit conversion for turnover rate constraints

In practice, fluxes are commonly expressed in mmol/gDCW/h, concentrations in mol/L, and TR constraints in units of inverse time. To make units consistent, we may need to scale the TR constraint using appropriate conversion factors, including the dry-cell weight-to-wet-cell weight ratio (gDCW/gWCW) and an approximate cell density (gWCW/L).

### TR-aware samplers

Sampling algorithms such as ACHR^55^ and optGpSampler^56^ are designed to operate on convex feasible sets. Consequently, they cannot be applied directly to mixed-integer formulations that include binary variables, such as models with thermodynamic constraints coupled to concentrations through linearization, or TR constraints introduced via binning.

A common workaround is to sample within a single flux directionality profile (FDP) ^49^. Fixing all reversible reactions to a chosen direction removes the need for the associated binary variables, enabling convex sampling. This approach also removes the coupling between the flux and concentration spaces created by thermodynamic constraints, because the linking binary variables can be dropped only after directionality is fixed.

For TR constraints, assigning an FDP is not sufficient. Even within a single FDP, the TR constraints still rely on piecewise or binned linearization that introduces binary variables. These variables cannot be removed without eliminating the approximation linking the consumption flux term to the concentration variable.

To address this, we modified the sampling procedure to generate only steady-state samples from regions that satisfy the TR constraints, while maintaining convex sampling steps wherever possible. We first illustrate the idea with a one-dimensional example, then generalize it to ACHR and optGpSampler steps.

The sampling procedures described below were developed for lower-bound turnover constraints, but their applicability does not generally extend to upper-bound constraints.

#### Example of sampling on a line

Given two points *x* (previous sample) and *wp* (warm-up point), we draw a candidate point *p* on the line segment defined by these points and restricted by the bounds of the convex feasible set,

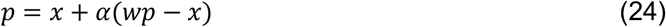

where *α* is constrained by feasibility in the convex space. Assuming a metabolite subject to a turnover rate lower bound 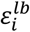, with 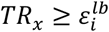 and 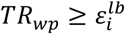, the following relations hold

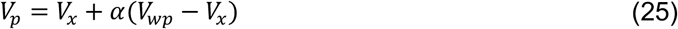

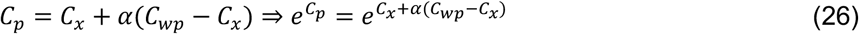

Therefore, the turnover rate of the metabolite at point *p* is

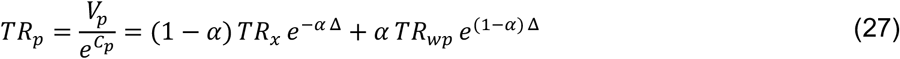

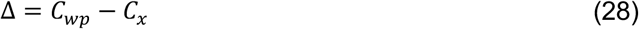

For *α* ∈ [0,1], *TR*_*p*_ will always satisfy the lower bound constraint. For values of *α* smaller than 0 or larger than 1, this does not generalize and will depend on the specific values of *TR*_*x*_,*TR*_*wp*_ and Δ. The range of allowed *α* is highly dependent on |Δ|, and as Δ approaches 0, the range becomes larger.

#### Sampling in a direction

In ACHR and in each Markov chain of optGpSampler, a sampling step selects a direction defined by the vector (*wp* − *cp*) and proposes a new point *p* starting from the current sample *x*

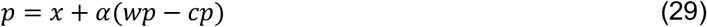

where the step size *α* is chosen within a feasibility interval determined by the convex constraints. Like in the previous example, the turnover rate at the proposed point can be written as

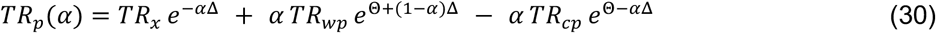

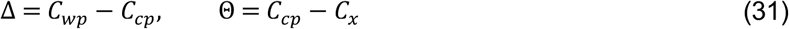

This creates a much more complex function, *TR*_*p*_(*α*) which can intersect the TR constraint lower bound *ε* at most twice. We ensure TR feasibility during sampling using the following procedure:

1. Determine the feasible interval for *α* imposed by the convex constraints (the standard ACHR bound computation).
2. Within this interval, locate up to two roots of *TR*_*p*_(*α*) − *ε* = 0. We bracket candidate roots by partitioning the interval and searching for sign changes, then apply the Brent method to identify the roots.
3. Restrict sampling of *α* to the sub-intervals where *TR*_*p*_ ≥ *ε*. Because *V*(*α*) − *ε exp*(*C*(*α*)) is concave, these admissible regions are contiguous and well defined.

#### Sampling an initial set of warmup points

We generate warmup points directly by solving the MILP formulation, ensuring that each warmup point satisfies the TR constraints. These points are also feasible for the corresponding LP relaxation. We encourage generating more warmup points than the conventional heuristic of 2x(number of variables) to provide a larger set of feasible directions for exploration. In this study, we generated 3x(number of variables) warmup points and a thinning step of 100 points.

#### Calculating the initial center point

Assume we have *k* warmup points *wp*_*i*_, each with values 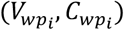. The initial center point is defined as the mean of the warmup points

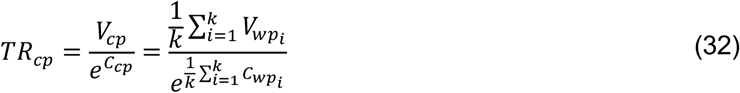

Using 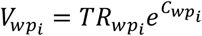, we obtain

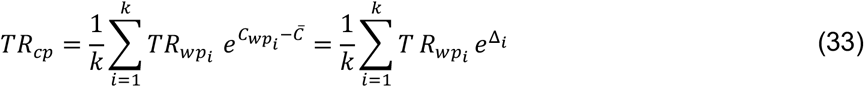

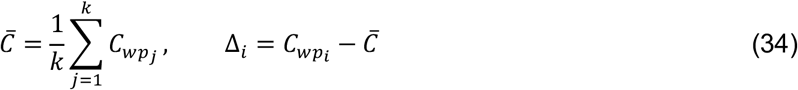

If each warmup point satisfies the turnover constraint 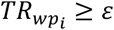 then it can be shown that this holds true for the center point too, as follows:

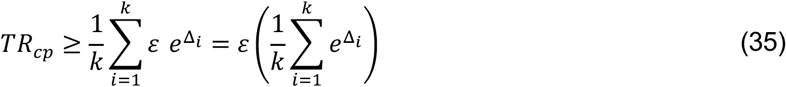

By Jensen’s inequality, since *e*^*x*^ is convex,

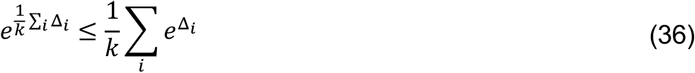

But 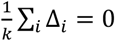 by definition, so

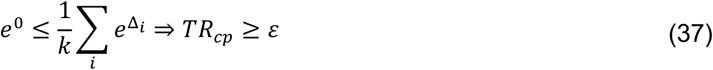

The average center point (*V*_*cp*_, *C*_*cp*_) will always satisfy the turnover constraint *TR*_*cp*_ ≥ *ε* whenever each warmup point individually satisfies 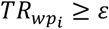.

#### Recalculation of the center point

After each sampling step, the center point is updated recursively as:

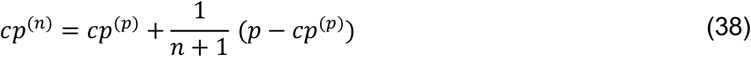

where *cp*^(*p*)^ is the previous center point (before adding the *n*-th sample), *p* is the newly generated sample, and *n* is the number of samples produced so far. This update can be seen as a convex combination of the previous center point and the new sample. It is analogous to the formulation of sampling along a line:

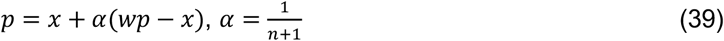

Since *n* ≥ 0 it follows that *α* ∈ [0,1], and in this range, the turnover ratio of the interpolated point satisfies 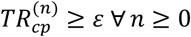.

### Imposing thermodynamic constraints

Thermodynamics Flux Analysis (TFA) ^11^ represents each reversible reaction by split, non-negative, forward and backward fluxes, 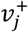and 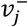. The binary variables 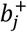 and 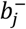 indicate whether the forward or backward direction is thermodynamically feasible. To impose thermodynamically feasible directionality, TFA uses equations (5)-(7) and the following constraints:

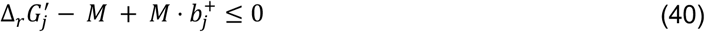

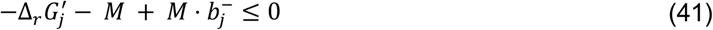

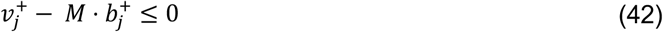

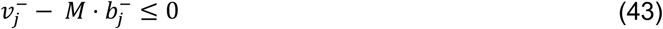

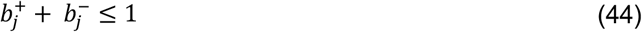

where *M* is a sufficiently large constant.

### Models used in this study

#### Reduced glycolysis

The reduced glycolysis model was generated from the known biochemistry of the *E. coli* glycolytic pathway. A complete schematic is shown in Figure 2A along with the names and symbols of reactions and metabolites. We do not model metabolites that are not directly related to carbon flow, and we assume that the reactions operate only in the direction of producing metabolites pyr and accoa. We lumped all the steps from g6p to pep into a single, kinetically reversible reaction LmpdG (*v*_2_). The remaining reactions exhibit kinetically irreversible rate laws (Supplementary Note 2). The glucose concentration is treated as a parameter and set to 10 g/L, with an uptake rate of 5 mmol gDCW^-1^ h^-1^.

#### Anaerobic *E. coli*

We used an existing, previously published kinetic model of anaerobic *E. coli* central carbon metabolism ^25^ and briefly summarize its construction here. The model is based on a reduced stoichiometric network generated with redGEM ^57^ and lumpGEM ^58^, covering core pathways and a lumped growth reaction. Anaerobic conditions were enforced by blocking oxygen uptake and oxygen-dependent respiratory reactions. The resulting steady-state model comprised 180 reactions and 141 metabolites.

To obtain a context-specific steady-state reference ensemble, we imposed physiological and experimental constraints ^26^, including fixed growth and glucose uptake rates, bounds on extracellular exchange consistent with the reported medium and secretions, and thermodynamic feasibility constraints. We then sampled thermodynamically feasible steady-state profiles using pyTFA, each including fluxes, metabolite concentrations, and thermodynamic variables. The kinetic model structure was generated using SKiMpy ^59^ and comprises 63 mass balances and 138 reaction rate laws spanning six mechanism families (Convenience, Generalized Reversible Hill, Bi-Uni Reversible Hill, Reversible Michaelis-Menten, Uni-Bi Reversible Hill, and a multi-substrate Monod form for growth), parameterized by 478 kinetic parameters.

#### Aerobic *E. coli*

For the experimental data-integration case study, we used the published *E. coli* core model of central carbon metabolism^32^. We extended the network with the Entner–Doudoroff pathway by adding phosphogluconate dehydratase (EDD), KDPG aldolase (EDA), and the intermediate KDPG. After removing blocked reactions and disconnected metabolites, the model comprised 89 reactions and 69 metabolites. We imposed thermodynamic constraints using pyTFA and fixed growth at 0.77 h^-1^ under aerobic conditions. The corresponding kinetic model was generated using SKiMpy.

#### Human cancer

We used a previously published reduced ovarian cancer metabolic model with integrated multi-omics constraints ^27^. Briefly, the model was derived from Recon3D and contextualized for *BRCA1*^WT^ and *BRCA1*^MUT^ ovarian cancer cells. In this study, we used the model representing the *BRCA1*^WT^ physiology with 2,359 reactions and 974 metabolites. The kinetic scaffold was built using SKiMpy and contains 716 mass balances (along with 25 pool-conservation equations) and 2,131 rate laws, requiring parametrization for 7,463 K_M_s and 2,131 V_max_s.

### Parametrization and linear dynamics

Several approaches can assign kinetic parameter values using information from a reference steady state, including methods such as ORACLE, GRASP, and closed-form parameterizations. Here, we used the ORACLE framework, which samples enzyme saturations around the reference steady-state metabolite concentrations, maps these saturations to Michaelis constants (K_M_s), and then back-calculates maximum velocities (v_max_s) to match the reference fluxes.

For each parameterized model, we computed the Jacobian matrix (Supplementary Note 1) and its eigenvalues to assess local stability and implied linear timescales. We retained only stable models, which had all eigenvalues with negative real parts. We also pruned them based on relevant linear timescales, estimated from the absolute value of the maximal eigenvalue. We define the characteristic time of the model as

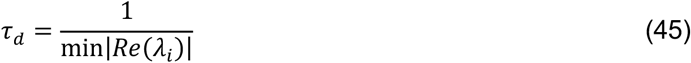

where *λ*_*i*_ are the eigenvalues of the Jacobian. The characteristic time defines the system’s slowest timescale and is associated with the TR values of the steady state sample (Supplementary Note 1). This definition was used to calculate the characteristic relaxation times reported in Figures 2-6.

### Modal analysis

To characterize the local dynamics around a steady state, we performed modal analysis of the nonlinear kinetic system

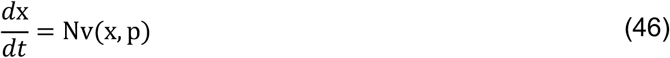

For a steady state solution x^*^ (Nv(x^*^, p) = 0), we define concentration perturbations as *δ*x(*t*) = x(*t*) − x^*^ and we linearize around x^*^

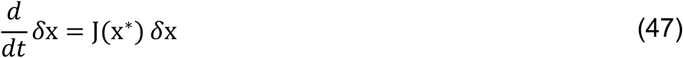

If the Jacobian is diagonalizable, we have:

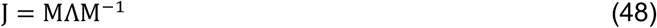

where Λ = diag(*λ*_1_, …, *λ*_*N*_) contains the eigenvalues *λ*_*i*_, and the columns of M are the corresponding right eigenvectors. Substituting this decomposition into the linearized system and introducing the modal mapping y = M^−1^*δ*x gives:

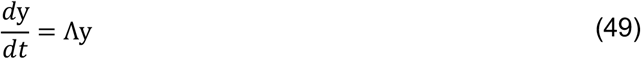

which decouples into independent scalar equations where each mode evolves as:

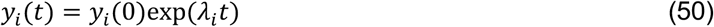

Thus, the local relaxation dynamics decompose into independent dynamic modes, each characterized by a specific eigenvalue *λ*_*i*_. The real part of *λ*_*i*_ defines the associated timescale 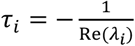. Each modal variable *y*_*i*_ represents a weighted combination of metabolite perturbations

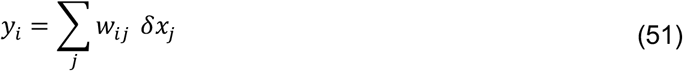

where *w*_*ij*_ denote the modal weights obtained from M^−1^. These weights quantify the contribution of metabolite *j* to mode *i*. Consequently, groups of metabolites evolve collectively on the timescale defined by the eigenvalue associated with that mode. To assess dynamical performance, we identified eigenvalues whose associated timescales *τ*_*i*_ were slower than a designated threshold. For these slow modes, metabolites with large modal weights |*w*_*ij*_| were interpreted as the primary contributors to the limiting dynamics. This procedure enabled the systematic identification of metabolites governing slow relaxation behavior in the kinetic models.

To ensure that we identified structurally limiting metabolite pools rather than artifacts of a single steady state or parameter set, we repeated this procedure across multiple steady states and independently sampled kinetic parameter sets. Aggregating modal contributions across this ensemble enabled the identification of globally persistent slow modes and metabolite pools that consistently govern the network’s limiting dynamic behavior.

### Iterative modal analysis workflow

To identify and reduce dynamically slow modes in the ovarian cancer kinetic model, we implemented an iterative modal analysis procedure coupled to targeted TR tightening. The goal was to selectively constrain metabolites responsible for undesired slow linear timescales while preserving maximal flexibility elsewhere.

#### Initial kinetic-model population

We first generated an unbiased population of kinetic models around the sampled steady states. Specifically, we selected 100 steady-state profiles and, for each, generated 100 independent parameterizations using ORACLE, yielding 10,000 kinetic models. For each model, we computed the Jacobian matrix and its eigenvalues and eigenvectors to characterize local linear dynamics.

#### Identification of slow modes

We defined slow modes as those associated with eigenvalues with a real part greater than −0.1 h^-1^. This cutoff was chosen relative to the physiological growth-rate scale and served to identify modes operating on timescales incompatible with the intended dynamical regime. For each flagged mode, we examined the corresponding eigenvector and identified metabolites with absolute modal weight greater than 0.1. The union of these metabolites across the population defined the set of dynamically influential species. In addition, all metabolites belonging to strictly linear pathway segments (i.e., only one consuming and one producing reaction) were included, because such metabolites are directly linked to an eigenvalue of the system, as shown in Supplementary Note 2.

#### Targeted TR tightening

We imposed a TR cutoff corresponding to a timescale 15-fold faster than the doubling time of the cell on the identified metabolite set and generated a new population of steady state samples. A new kinetic-model population was then generated and re-evaluated via modal analysis.

#### Iterative procedure

The above procedure was repeated until the sampled kinetic-model population achieved the targeted linear dynamics, or until the set of metabolites associated with slow modes changed only marginally between successive iterations. In practice, three iterations removed most slow modes while preserving feasible steady-state variability. After convergence, optional global tightening of the TR cutoff was explored to further shift the characteristic time distribution toward faster timescales.

### Nonlinear simulations

#### Bioreactor simulations

We simulated batch fermentation of anaerobically grown *E. coli* using the bioreactor framework described in our prior work ^25^. Each intracellular kinetic model was extended with standard bioreactor mass-balance equations, including ODEs for biomass growth (biomass concentration) and extracellular metabolite dynamics based on uptake and secretion fluxes. Simulations were initialized with the steady-state concentrations and fluxes used for parameterization, together with the inoculum size and medium composition reported by Varma and Palsson ^26^.

#### *In silico* drug administration simulations

We performed simulations using a procedure described in our prior work ^27^. Briefly, we modeled drug action as a first-order process that reduces the target enzyme activity *E*_tot_ toward a prescribed fractional level (TMDS enzyme; 10-fold reduction over an 8-hour window). For the simulation, we used (i) all kinetic models obtained after the third iteration of the iterative workflow that satisfied the characteristic-time pruning criterion (169) and (ii) a representative subset of the initial unconstrained ensemble (150 models selected so that their characteristic times were evenly spaced across the original distribution). Each model was integrated from its reference steady state until it reached a new steady state or 600 simulation hours, whichever occurred first. We quantified the post-perturbation response from the growth-rate trajectory and the time required to reach a new steady value.

### Equilibrium detection algorithm

To quantify the time required for the perturbed system to reach a new operating point, we implemented an equilibrium (steady state) detection algorithm. A post-perturbation equilibrium corresponds to any state *x*^*^ satisfying:

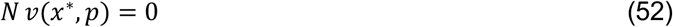

Because numerical trajectories never yield an exact zero derivative, we formulated equilibrium detection using scaled, persistent criteria evaluated over a time window of length Δ. At time *t*, we define the residual vector *r*(*t*) = *Nv*(*x*(*t*), *p*) and the scaled residual norm:

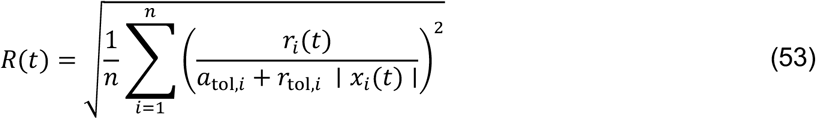

where *a*_*tol,i*_ and *r*_*tol,i*_ are metabolite-specific absolute and relative scaling parameters. This norm measures whether the instantaneous rates of change are small compared to the scale of each metabolite concentration.

To avoid falsely identifying equilibrium when the trajectory only briefly passes near a slow manifold, we require that the residual remain small continuously over the entire time window:

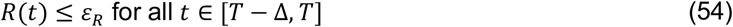

where *T* is the current time, and *ε*_*R*_ is a global threshold.

Residual smallness alone can be misleading because cancellations in *Nv*(*x, p*) may produce small derivatives even while the state drifts slowly. We therefore impose a mandatory drift criterion over the same time window. We define the scaled state change:

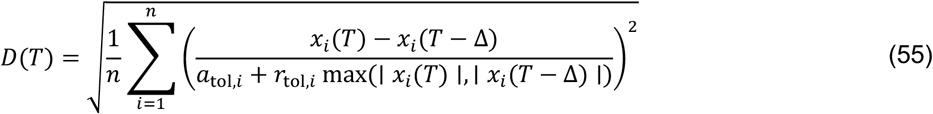

and the corresponding drift rate 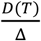. We assume that the system has reached equilibrium at time *T* if and only if both conditions are satisfied:

i. Residual persistence holds over [*T* − Δ, *T*].
ii. 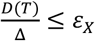 where *ε*_*X*_ controls the maximum allowed scaled rate of change over the window.

The algorithm determines whether the system has stopped evolving, regardless of the specific equilibrium reached. The equilibration time identified by this procedure is governed by the slowest stable dynamical mode of the perturbed system. Because very slow models can exhibit small instantaneous derivatives while still drifting gradually, we imposed a strict drift threshold to avoid falsely declaring equilibrium in such cases. For this study, we used *a*_*tol*_ = *r*_*tol*_ = 10^−6^ for all metabolites, a time window of 10 hours, and *ε*_*R*_ = *ε*_*X*_ = 1.

### Derivation of literature-based turnover ranges

We used data from the glucose-grown cultures reported in Gerosa et al. ^30^ and Park et al. ^29^, the uninduced wild-type controls from the catabolic and anabolic experimental series of Kochanowski et al. ^31^, and the aerobic glucose-limited chemostat cultures of Taymaz-Nikerel et al. ^28^. We mapped these data to the published *E. coli* core model^32^, extended here with the Entner–Doudoroff pathway, and retained only metabolites with measured concentrations and flux estimates for all connected model reactions. For each metabolite, we calculated the turnover rate by dividing the total stoichiometry-weighted consuming flux by the corresponding metabolite pool, after converting all quantities to consistent units. We calculated turnover time as the reciprocal of the turnover rate. For Taymaz-Nikerel et al., we used the reported turnover times as central estimates and propagated one standard error in the corresponding metabolite concentrations to obtain interval bounds.

For the remaining studies, intervals were constructed using the uncertainty measures reported in each source: one standard deviation above and below the mean for Gerosa et al. and Kochanowski et al., and the reported 95% bounds for Park et al. Where multiple estimates were available for the same metabolite, we defined the combined interval using the lowest lower bound and the highest upper bound. This procedure yielded intervals for 18 metabolites that incorporate both reported experimental uncertainty and variation among experimental conditions. The final intervals are provided in Supplementary Table S2.

## Data Availability

The data generated in this study have been deposited in the Zenodo repository https://doi.org/10.5281/zenodo.21133382.

## Code Availability

The code used to develop the model, perform the analyses, and generate results in this study is publicly available and has been deposited in https://github.com/EPFL-LCSB/METEOR-K under the Apache-2.0 license. The kinetic models were built using SKiMPy (Symbolic Kinetic Models in Python), available at https://github.com/EPFL-LCSB/skimpy.

## Acknowledgements

This work was supported by funding from the Swiss National Science Foundation grants 200021_188623 and CRSII5_198543, the European Union’s Horizon 2020 research and innovation programme under grant agreement 814408, and the École Polytechnique Fédérale de Lausanne (EPFL).

## Author Contributions

I.T., D.W., G.F, and L.M. conceptualized the overall methodology, V.H. and L.M. acquired funding and supervised the research, I.T. curated the data, models, and computational resources, I.T., D.W., B.N., and G.F. designed the code, I.T., D.W., G.F, B.N., analyzed the data, I.T. validated results, I.T. and L.M. visualized the results, I.T. prepared the original draft of the manuscript, I.T., B.N., and L.M. reviewed and edited the draft. All authors read and commented on the manuscript.

## Declaration of generative AI and AI-assisted technologies in the writing process

During the preparation of this work, the authors used ChatGPT to improve the manuscript’s readability and language. After using this tool, the authors reviewed and edited the content as needed and take full responsibility for the content of the published article.

