## Supplementary information for "Scalable biophysical constraints link steady-state and kinetic metabolic modeling"

Toumpe I. et al

#### Supplementary Note 1

**Analytical expression of eigenvalues in metabolic networks**

The Jacobian of a biochemical reaction network can be written in factored matrix form (Wang et al., 2004) as:

$$J = N \cdot V \cdot E \cdot X^{-1}$$

Here, $N$ is the stoichiometric matrix (metabolites x reactions), $V$ is a diagonal matrix of steady-state reaction fluxes, $E$ is the elasticity matrix with respect to metabolites, and $X$ is a diagonal matrix of steady-state metabolite concentrations. With these definitions, each Jacobian entry $a_{kl}$ takes the form:

$$a_{kl} = \sum_{j=1}^{n_r} \eta_{kj} \frac{v_j}{x_l} \varepsilon_{x_l}^{v_j}$$

Where $\eta_{kj}$ is the stoichiometric coefficient of metabolite $k$ in reaction $j$, $v_j$ is the steady-state flux of reaction $j$, $x_l$ is the steady-state concentration of metabolite $l$, $\varepsilon_{x_l}^{v_j}$ is the elasticity of $v_j$ with respect to $x_l$ and $n_r$ is the number of reactions.

Let $M$ and $M^{-1}$ denote the matrices that diagonalize $J$ so that $J = M \cdot \Lambda \cdot M^{-1}$. Let $m_{li}$ and $n_{ik}$ be the elements of $M$ and $M^{-1}$, respectively. Then the eigenvalues, the diagonal entries of $\Lambda$, can be expressed as:

$$\lambda_{ii} = \sum_{j=1}^{n_r} \sum_{k=1}^{n_m} \sum_{l=1}^{n_m} m_{li} n_{ik} \eta_{kj} \frac{v_j}{x_l} \varepsilon_{x_l}^{v_j}$$

where $n_m$ is the number of metabolites. Therefore, each eigenvalue is a weighted linear combination of terms of the form $\frac{v_j}{x_l} \varepsilon_{x_l}^{v_j}$, that is, elasticity terms scaled by the corresponding flux-to-concentration ratios. In sparsely connected regions of a network, only a small number of these terms may contribute to a given eigenvalue. An explicit example is provided for the reduced glycolysis model in Supplementary Note 2.

#### Supplementary Note 2

**Structure of the reduced glycolysis model**

The reduced glycolysis model can be described by the following system of ordinary differential equations:

$$\frac{d}{dt}\begin{bmatrix} X \\ Y \\ Z \\ T \end{bmatrix} = \begin{bmatrix} 1 & -1 & 0 & 0 & 0 & 0 \\ -1 & 2 & -1 & 0 & 0 & 0 \\ 1 & 0 & 1 & -1 & -1 & 0 \\ 0 & 0 & 0 & 0 & 1 & -1 \end{bmatrix} \begin{bmatrix} v_1(V_{max1}, S, K_{m,S}, Y, K_{m,Y1}) \\ v_2(V_{max2}, X, K_{m,X}, Y, K_{m,Y2}, K_{eq,2}) \\ v_3(V_{max3}, Y, K_{m,Y3}) \\ v_4(V_{max4}, Z, K_{m,Z4}) \\ v_5(V_{max5}, Z, K_{m,Z5}) \\ v_6(V_{max6}, T, K_{m,T}) \end{bmatrix}$$

The symbols of metabolites and fluxes are denoted in Figure 2A. The reaction rates are defined as follows:

Reaction one, glucose transport via pep-pyr phosphotransferase system (GLCpts), follows irreversible Bi-Bi kinetics:

$$v_1 = V_{max1} \frac{\frac{S}{K_{m,S}} \frac{Y}{K_{m,Y1}}}{\left(1 + \frac{S}{K_{m,S}}\right)\left(1 + \frac{Y}{K_{m,Y1}}\right)}$$

Reaction two, lumped glycolysis (LmpdG), follows reversible uni-bi kinetics:

$$v_2 = V_{max2} \frac{\frac{X}{K_{m,X}}\left(1 - \frac{1}{K_{eq,2}} \frac{Y^2}{X}\right)}{1 + \frac{X}{K_{m,X}} + 2\frac{Y}{K_{m,Y2}} + \left(\frac{Y}{K_{m,Y2}}\right)^2}$$

Reaction three, pyruvate kinase (PYK), follows 4<sup>th</sup>-order irreversible Hill kinetics:

$$v_3 = V_{max3} \frac{\frac{Y^4}{K_{m,Y3}{}^4}}{\left(1 + \frac{Y^4}{K_{m,Y3}{}^4}\right)}$$

Reaction four, lactate dehydrogenase (LDH), follows irreversible Michaelis-Menten kinetics:

$$v_4 = V_{max4} \frac{\frac{Z}{K_{m,Z4}}}{\left(1 + \frac{Z}{K_{m,Z4}}\right)}$$

Reaction five, pyruvate dehydrogenase (PDH), follows irreversible Michaelis-Menten kinetics:

$$v_5 = V_{max5} \frac{\frac{Z}{K_{m,Z5}}}{\left(1 + \frac{Z}{K_{m,Z5}}\right)}$$

Reaction six, acetaldehyde dehydrogenase (ACALD), follows irreversible Michaelis-Menten kinetics:

$$v_6 = V_{max6} \frac{\frac{T}{K_{m,T}}}{\left(1 + \frac{T}{K_{m,T}}\right)}$$

**Analytical Jacobian structure and eigenvalue decomposition**

The Jacobian matrix $J$ allows us to analyze the dynamic behavior of a system around its steady state $x_{ss}$. If we define the deviation $\tilde{x}(t)$ from steady state $x_{ss}$ as $\tilde{x}(t) = x(t) - x_{ss}$, then we can describe the linearized system around $x_{ss}$ as:

$$\frac{d\tilde{x}}{dt} = \mathrm{J}\,\tilde{x}$$

The Jacobian matrix of the studied model has the following form:

$$J = \begin{bmatrix} -\frac{\partial v_2}{\partial X} & \frac{\partial v_1}{\partial Y} - \frac{\partial v_2}{\partial Y} & 0 & 0 \\ 2\frac{\partial v_2}{\partial X} & 2\frac{\partial v_2}{\partial Y} - \frac{\partial v_1}{\partial Y} - \frac{\partial v_3}{\partial Y} & 0 & 0 \\ 0 & \frac{\partial v_1}{\partial Y} + \frac{\partial v_3}{\partial Y} & -\frac{\partial v_4}{\partial Z} - \frac{\partial v_5}{\partial Z} & 0 \\ 0 & 0 & \frac{\partial v_5}{\partial Z} & -\frac{\partial v_6}{\partial T} \end{bmatrix}$$

The eigenvalues of the Jacobian matrix can be computed as roots of the equation $\det(\lambda I - J) = 0$. From the above matrix, we obtain the characteristic equation:

$$\left[\left(\lambda + \frac{\partial v_2}{\partial X}\right)\left(\lambda - 2\frac{\partial v_2}{\partial Y} + \frac{\partial v_1}{\partial Y} + \frac{\partial v_3}{\partial Y}\right) - 2\frac{\partial v_2}{\partial X}\left(\frac{\partial v_1}{\partial Y} - \frac{\partial v_2}{\partial Y}\right)\right]\left(\lambda + \frac{\partial v_4}{\partial Z} + \frac{\partial v_5}{\partial Z}\right)\left(\lambda + \frac{\partial v_6}{\partial T}\right) = 0$$

The magnitude of the eigenvalues is dictated by the elements of the Jacobian. As was shown in Supplementary Note 1, the elements of the Jacobian are a function of an elasticity term scaled by the corresponding flux-to-concentration ratio.

The Jacobian of the reduced glycolysis model has a block-lower-triangular structure, consisting of a coupled 2×2 block for $X$ and $Y$, followed by scalar diagonal blocks associated with $Z$ and $T$. Although perturbations can propagate from $Y$ to $Z$ and from $Z$ to $T$, neither $Z$ nor $T$ feeds back into the upstream blocks. Consequently, the diagonal entries associated with $Z$ and $T$ are exact eigenvalues. In particular, accoa ($T$; Figure 2A of the main manuscript) is produced by pyruvate dehydrogenase (PDH, $v_5$) and consumed by acetaldehyde dehydrogenase (ACALD, $v_6$). Because $v_6$ is the only reaction rate that depends on $T$, the last two factors of the characteristic equation yield:

$$\lambda_Z = -\left(\frac{\partial v_4}{\partial Z} + \frac{\partial v_5}{\partial Z}\right), \qquad \lambda_T = -\frac{\partial v_6}{\partial T}$$

For the irreversible rate kinetics of $v_6$

$$\lambda_T = -\frac{v_6}{T}\varepsilon_T^{v_6} = -\mathrm{TR}_T\varepsilon_T^{v_6}$$

with

$$\varepsilon_T^{v_6} = \frac{1}{1 + \frac{T}{K_{m,T}}}, \quad 0 < \varepsilon_T^{v_6} \leq 1$$

Therefore, $|\lambda_T| \leq TR_T$, or, equivalently, the accoa relaxation time satisfies $\tau_T = \frac{1}{|\lambda_T|} \geq \frac{1}{TR_T}$. Because the characteristic relaxation time of the complete system is determined by its slowest mode, it cannot be shorter than $\tau_T$. Thus, when accoa is the limiting mode, $\frac{1}{TR_T}$ defines the fastest attainable characteristic time, while the sampled elasticity determines how closely an individual model approaches this limit. If another mode is slower, that mode instead determines the characteristic relaxation time, as observed when the turnover of accoa is increased in Figure 2D.

This structure commonly occurs in feed-forward linear pathway segments. A downstream metabolite may receive perturbations from upstream variables, producing entries below the block diagonal, while not feeding back into the upstream reaction rates. When this structure produces a scalar diagonal block, its diagonal entry is an exact eigenvalue. For an irreversible Michaelis–Menten consuming reaction, this eigenvalue can be expressed as the product of the metabolite turnover rate and the corresponding elasticity.

This does not necessarily hold when metabolites are part of branched or cyclic pathway structures. The Jacobian contains larger coupled blocks (for example, the 2×2 block in $X$ and $Y$). In these cases, eigenvalues depend on both diagonal and off-diagonal terms, so their magnitudes are not determined by a single turnover rate and elasticity. Nevertheless, if the Jacobian is diagonally dominant, so that the off-diagonal elements are small relative to the diagonal entries, then the diagonal terms largely determine the eigenvalue range. In this case, turnover rates provide a practical approximation to the expected eigenvalue magnitudes, even in the presence of weak coupling.

**Effect of reaction reversibility near equilibrium**

To examine the effect of reaction reversibility on the eigenvalues of the system, we assume that acetaldehyde dehydrogenase (ACALD, $v_6$) follows reversible uni-uni Michaelis-Menten kinetics. Let P denote the product concentration and assume that P is held fixed. The reversible rate law is:

$$v_6^{\text{rev}} = \frac{V_{max,6}^{+}\,\dfrac{T}{K_{m,T}} - V_{max,6}^{-}\,\dfrac{P}{K_{m,P}}}{1 + \dfrac{T}{K_{m,T}} + \dfrac{P}{K_{m,P}}} = \frac{V_{max,6}^{+}\,\dfrac{T}{K_{m,T}}(1 - \Gamma_6)}{1 + \dfrac{T}{K_{m,T}} + \dfrac{P}{K_{m,P}}}$$

where the displacement from equilibrium is defined as:

$$\Gamma_6 = \frac{V_{max,6}^{-}\,\dfrac{P}{K_{m,P}}}{V_{max,6}^{+}\,\dfrac{T}{K_{m,T}}} = \frac{P}{K_{eq,6}T}$$

For the net-consuming direction considered here, $0 \leq \Gamma_6 < 1$, and $\Gamma_6 \to 1^{-}$ denotes approach to equilibrium. The elasticity of the reversible reaction with respect to accoa is then:

$$\varepsilon_T^{v_6,\text{rev}} = \frac{T}{v_6^{\text{rev}}}\frac{\partial v_6^{\text{rev}}}{\partial T} = \frac{1}{1 - \Gamma_6} - \frac{\dfrac{T}{K_{m,T}}}{1 + \dfrac{T}{K_{m,T}} + \dfrac{P}{K_{m,P}}}$$

For comparison, the irreversible elasticity is bounded between zero and one. In contrast, the first term in the reversible elasticity increases without bound as the displacement from equilibrium approaches one, whereas the saturation term remains finite. The reversible elasticity can consequently exceed one.

Because the product concentration is fixed, the reversible reaction remains the only reaction rate that depends on accoa. The block-lower-triangular structure of the Jacobian is therefore unchanged. Only the expression for its final diagonal element changes, and this element remains an exact eigenvalue:

$$\lambda_T = J_{TT}^{\mathrm{rev}} = -\frac{\partial v_6^{\mathrm{rev}}}{\partial T} = -\frac{v_6^{\mathrm{rev}}}{T}\varepsilon_T^{v_6,\mathrm{rev}} = -\mathrm{TR}_T\varepsilon_T^{v_6,\mathrm{rev}}$$

Holding the net steady-state flux and accoa concentration fixed also fixes the accoa turnover rate. Across kinetic parameterizations with different displacements from equilibrium, the normalized elasticity and hence the accoa eigenvalue can nevertheless change. Conversely, at fixed thermodynamic displacement and saturation state, changing the turnover rate scales the eigenvalue. Turnover, thermodynamic displacement, and kinetic saturation therefore act jointly.

As $\Gamma_6$ approaches one, the accoa eigenvalue becomes increasingly negative and the accoa mode becomes fast. It then ceases to determine the characteristic relaxation time of the complete system, which shifts to the eigenvalue associated with pyr or to the upstream 2×2 block in g6p and pep. These remaining modes are still scaled by their corresponding flux-to-concentration ratios, although the eigenvalues of the coupled upstream block depend on combinations of turnover ratios, elasticities, and off-diagonal interactions. Thus, near-equilibrium reversibility can produce a fast local accoa mode despite low net accoa turnover, but it does not remove turnover scaling from the characteristic dynamics of the complete system; instead, the rate-limiting mode shifts to another part of the network.

The implication for turnover-based filtering depends on the total net turnover of the metabolite rather than on the contribution of the near-equilibrium reaction alone. In a more general network, suppose that accoa is also consumed by an additional non-near-equilibrium pathway with positive net steady-state flux, $v_{\mathrm{slow}}$. If the turnover threshold is calibrated from a physiologically acceptable slower response such that

$$\frac{v_{\mathrm{slow}}}{\mathrm{T}} \geq \mathrm{TR}_{\mathrm{min}}$$

then for the unit stoichiometries considered here,

$$\frac{v_6^{\mathrm{rev}} + v_{\mathrm{slow}}}{\mathrm{T}} \geq \frac{v_{\mathrm{slow}}}{\mathrm{T}} \geq \mathrm{TR}_{\mathrm{min}}$$

The state therefore satisfies the accoa turnover constraint irrespective of the near-equilibrium contribution. For the mechanism considered here, a false exclusion by the accoa turnover constraint can occur only when the total net accoa turnover remains below the imposed cutoff even though the complete Jacobian yields physiologically acceptable relaxation owing to the large near-equilibrium elasticity. This inequality concerns the turnover-filtering criterion only; if the additional pathway depends on a dynamic metabolite concentration, its sensitivity also enters the Jacobian and the resulting modes must be evaluated jointly.

### Supplementary Note 3

**Fastest attainable local dynamics for the reduced glycolysis model**

In Figures 2D and 2E, we showed that once the turnover ratio of accoa is increased to match the other metabolites in the system (timescale of 2s), g6p becomes the new limiting metabolite. This observation motivated us to explore what is the fastest characteristic time the model can achieve. To estimate this upper bound, we computed, for each metabolite independently, the maximum attainable TR by (i) maximizing the absolute sum of all consuming fluxes (accounting for stoichiometric coefficients) and (ii) minimizing the log-concentration within its feasible range. We report these results as inverse TR values so they can be interpreted directly as characteristic timescales:

| Metabolite | Fastest attainable timescale (s) |
|---|---|
| accoa | 0.336 |
| g6p | 1.50 |
| pyr | 0.0006 |
| pep | 0.0012 |

Based on this analysis, g6p is indeed the limiting metabolite of the model. Even under its best-case TR value, the implied minimum timescale is 1.50s. We then considered an idealized scenario in which all metabolites are fixed at their best achievable TR values and used these targets to construct a steady-state solution. Around this steady state, we sampled a large ensemble of kinetic parameters (100,000 models) and found that the fastest attainable characteristic time across the ensemble was 1.57s.

### Supplementary Note 4

**Implications of initial biomass precursor concentrations on bioreactor simulations**

To simulate batch bioreactor dynamics, we initialized each model with an experimentally derived inoculum and intracellular metabolite concentrations taken from the steady-state sample used to build the corresponding kinetic model. A key determinant of early bioreactor behavior is the initial abundance of biomass precursors, defined here as metabolites that participate in the model's growth rate reaction. When these precursor concentrations start low, growth is delayed until the precursor pools replenish, producing an extended lag phase.

To quantify this effect, we generated populations of kinetic models from steady states that differed primarily in biomass precursor concentrations. We constructed three groups of steady states: (i) 100 samples with biomass precursor concentrations between the lower bound and 10% of their feasible range, (ii) 100 samples with concentrations between 45% and 55% of their feasible range, and (iii) 100 samples with concentrations between 70% and 100% of their feasible range. The feasible range is derived by identifying the lower and upper bounds on each meetable concentration, given all the constraints of the steady-state model. For each steady state, we built 100 kinetic models, kept only the stable ones (all eigenvalues must have a negative real part), and aggregated their bioreactor simulation outputs. Figure S2 shows that both the biomass trajectories and the biomass concentration at t = 16 h differ substantially across these three groups. These results indicate that, for bioreactor simulations, besides enforcing TR constraints, we must also ensure sufficiently high initial biomass precursor concentrations to allow the models to evolve in faster dynamic scales.

### Supplementary Note 5

**Accuracy versus number of bins in the discretization method**

Assume a maximum allowed error of $m_a$. In the worst-case scenario, the active auxiliary variable $AC_m$ lies on the lower bin bound $\mathrm{lb}_m$, while the true concentration associated with the log variable $C_i$ is at the upper bound, so $C_i = \ln(\mathrm{ub}_m)$. For this mismatch to matter, the approximate TR value should be exactly equal to $\varepsilon_i$. Under these conditions, enforcing sufficient linearization accuracy requires the ratio of bin bounds to satisfy:

$$\frac{\mathrm{ub}_m}{\mathrm{lb}_m} \le \frac{\varepsilon_i}{\varepsilon_i - m_a}, \qquad \forall m,\ \ 0 < m_a < \varepsilon_i$$

Equivalently, in logarithmic form, the bin width must satisfy:

$$\ln(\mathrm{ub}_m) - \ln(\mathrm{lb}_m) \ \le \ \ln\left(\frac{\varepsilon_i}{\varepsilon_i - m_a}\right)$$

This bound highlights the benefit of logarithmic binning relative to linear binning. For example, with a TR cutoff $\varepsilon_i = 10$ and maximum allowed error of $m_a = 1$, linear discretization with $N$ = 200 bins yields several bins whose ratios $\frac{\mathrm{ub}_m}{\mathrm{lb}_m}$ violate the above condition (Figure S3). In contrast, logarithmic discretization satisfies the condition for all bins with substantially fewer bins ($N$ = 105, Figure S3). Moreover, log discretization enforces a constant ratio $\frac{\mathrm{ub}_m}{\mathrm{lb}_m}$ across bins, which makes the approximation error uniform over the feasible concentration range.

In practice, we select the discretization as follows:

1. Choose the TR threshold $\varepsilon_i$,

2. specify the maximum deviation $m_a$,

3. determine the feasible concentration range for the metabolite, and

4. compute the minimum number of bins required under logarithmic discretization over the feasible bounds.

**Accuracy versus number of breakpoints in the linearization method**

We approximate the exponential function in an interval between two breakpoints $[x_1, x_2]$ with a linear function:

$$f(x) = e^x, \qquad f_a(x) = ax + b$$

where $a = \frac{e^{x_2} - e^{x_1}}{x_2 - x_1}$ and $b = \frac{x_2 e^{x_1} - x_1 e^{x_2}}{x_2 - x_1}$. To quantify approximation quality $[x_1, x_2]$, we define the pointwise error:

$$error(x) = f_a(x) - f(x) = ax + b - e^x$$

To find where the error is maximal, we can compute the derivative and find its root:

$$\frac{derror(x^*)}{dx} = a - e^{x^*} = 0 \quad \Rightarrow \quad x^* = \ln(a) = \ln\left(\frac{e^{x_2} - e^{x_1}}{x_2 - x_1}\right)$$

Since $e^x$ is strictly convex, the $f_a(x)$ line lies above the function on the open interval $(x_1, x_2)$. Therefore $error(x) \ge 0$ and the single critical point $x^*$ gives the global maximum. Thus, the maximum error is:

$$e_{\max} = a\ln(a) + b - a.$$

It is worth pointing out that since $f_a(x) \ge f(x)$, the linearized exponential systematically overestimates the true value. In the TR approximation, this produces a conservative constraint because the predicted TR is smaller than the true TR, so $\varepsilon \le TR_{pr} \le TR_{actual}$.

To visualize the error across the concentration range, we use a representative metabolite with concentrations from $10^{-6}$ to 0.05 M. The approximation and its error are shown in Figure S4. The error increases as the interval shifts toward larger x, motivating a nonuniform breakpoint placement with higher density near the upper bound (Figure S4).

Instead of placing breakpoints with uniform spacing in log-concentration, we use a warped grid that concentrates points near the upper bound. Let the total log-concentration range be $[C_{lb}, C_{ub}]$ and let $N$ denote the number of breakpoints. Define a normalized parameter grid:

$$t_k = \frac{k}{N-1}, \qquad k = 0,1,\dots,N-1$$

and apply a power-law transformation

$$\phi_\alpha(t) = t^\alpha, \qquad \alpha \in (0,1]$$

The breakpoints are then given by

$$C_k = C_{lb} + \phi_\alpha(t_k)\,(C_{ub} - C_{lb}) = C_{lb} + \left(\frac{k}{N-1}\right)^\alpha (C_{ub} - C_{lb}).$$

For $\alpha = 1$, this reduces to uniform spacing. For $\alpha < 1$, more breakpoints are placed near $C_{ub}$, where the linearization error of $e^{C_i}$ is larger.

### Supplementary Note 6

**Bounding-box estimate of feasible-space volume**

We estimated the reduction in feasible-space volume using a bounding-box approximation. For each model, thermodynamic variability analysis was used to obtain the minimum and maximum feasible values of each log-concentration and flux variable, both before and after applying TR constraints.

For each variable $i$, the feasible width was defined as $w_i = |x_i^{max} - x_i^{min}|$. The feasible space was then approximated as a high-dimensional rectangular box, with one side length per variable. The corresponding bounding-box volume is:

$$V_{box} = \prod_j w_j$$

It is important to note that this estimate is conservative. It treats each variable range as independent and therefore ignores correlations and constraints coupling variables within the feasible space. As a result, the estimated volume represents an upper bound on the true feasible-space volume, and the actual feasible volume is expected to be substantially smaller.

Because this product spans many dimensions, the calculation was performed in $ln$ space. The reported volume ratio was computed as:

$$\ln\left(\frac{V_{constrained}}{V_{original}}\right) = \sum_j \ln\left(w_j^{constrained}\right) - \sum_j \ln\left(w_j^{original}\right)$$

Variables with zero feasible width in either condition were excluded from the calculation. The reported ratio, therefore, compares the active positive-width dimensions shared between the original and TR-constrained models. Because this estimate aggregates many dimensions and ignores correlations between variables, it should be interpreted as a high-dimensional search-space contraction induced by the targeted constraints, not as a per-variable uncertainty reduction.

**Summary of bounding-box volume estimates for the feasible space before and after TR constraints.**

| Model | Dimensions (non-zero width) | Log volume ratio | Raw Volume Ratio | Geometric-mean width ratio |
|---|---|---|---|---|
| Reduced glycolysis pathway (global TR enforcement equivalent to 5 seconds) | 16 (15) | -0.850 | $4.27*10^{-1}$ | 0.945 |
| Anaerobic *E. coli* (global TR enforcement: 10-fold faster than doubling time) | 313 (308) | -2.889 | $5.56*10^{-2}$ | 0.991 |
| Ovarian cancer (selective TR enforcement: 15-fold faster than doubling time applied for 705 metabolites) | 3329 (3315) | -21.789 | $3.44*10^{-10}$ | 0.993 |

Throughout the models, the TR constraints have a bigger effect on the feasible range of the concentration variables. As an example, we showcase the width ratios of the flux variables and concentration variables for the anaerobic *E. coli* model (Figure S6). Effectively, none of the flux variables had a non-negligible reduction in their feasible range, whereas some of the concentration variables had considerable reductions in their feasible space. Examples include nucleotide GTP (ratio of 0.31, participates in 3 reactions) and Glycerate (ratio of 0.56, participates in 4 reactions).

### Supplementary Figures and Tables

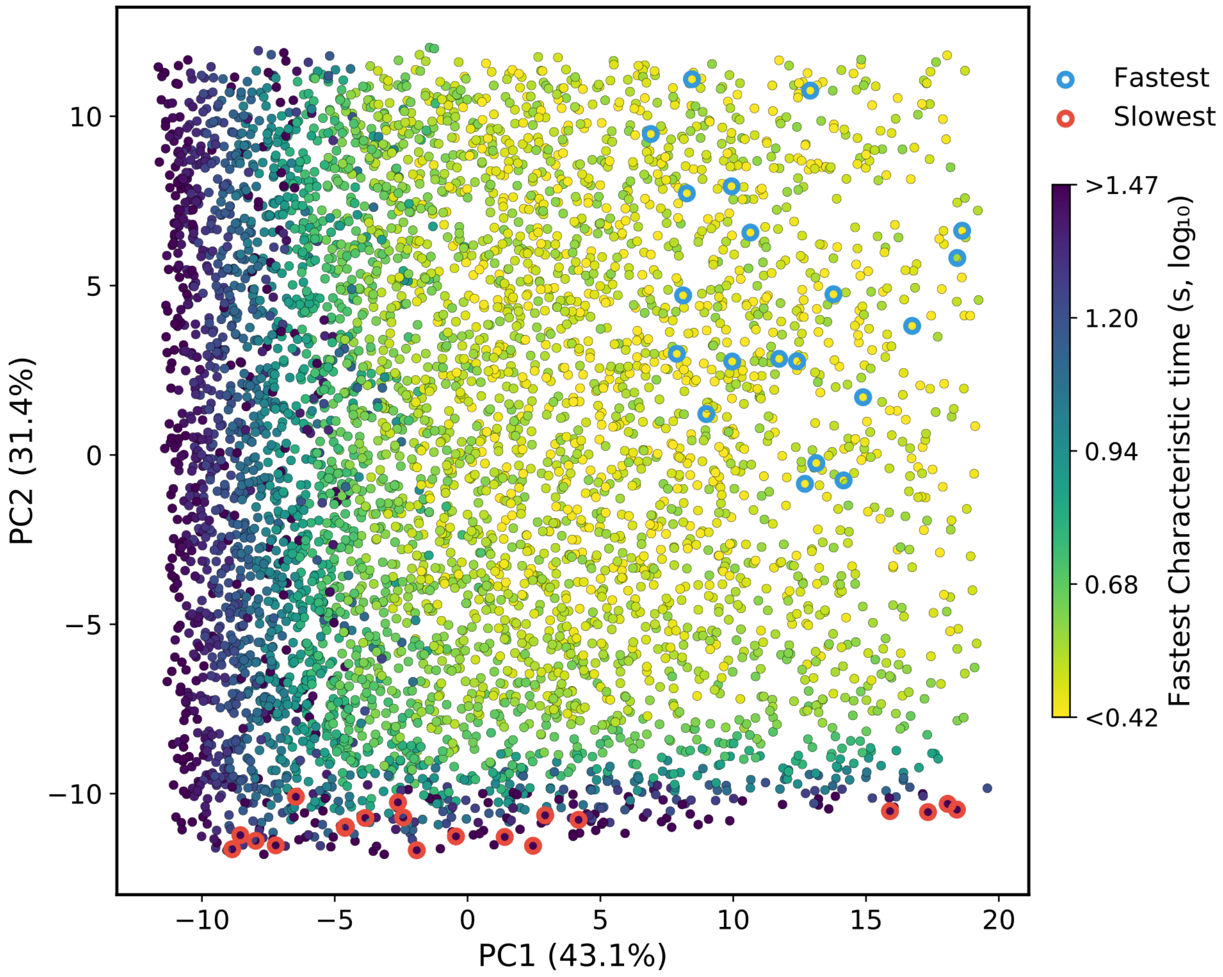


**Figure S1.** Principal component analysis (PCA) of 5,000 feasible steady state samples. Points are colored by the fastest characteristic time (log10 scale) computed from 100 kinetic models parameterized around each steady state. All characteristic times below the 5th percentile and above the 95th percentile of the distribution were clipped to the same color limits. Blue circles mark the 20 fastest steady states, and red circles mark the 20 slowest steady states by fastest characteristic time.

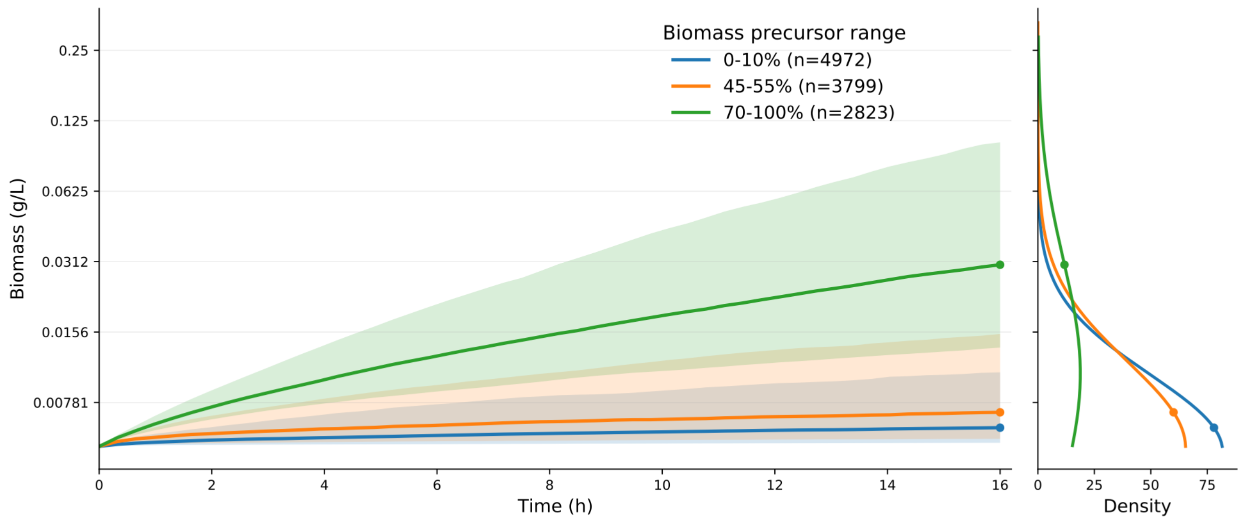


**Figure S2.** Initial biomass precursor abundance shapes biomass dynamics in batch bioreactors. Biomass trajectories (median with interquartile range) are shown for models initialized from steady states grouped by biomass precursor concentrations (0-10%, 45-55%, 70-100% of each precursor's feasible range). The right panel shows the corresponding biomass concentration distribution at $t = 16$h (log-normal fit). Higher initial precursor levels yield faster biomass accumulation and higher biomass at 16 h.

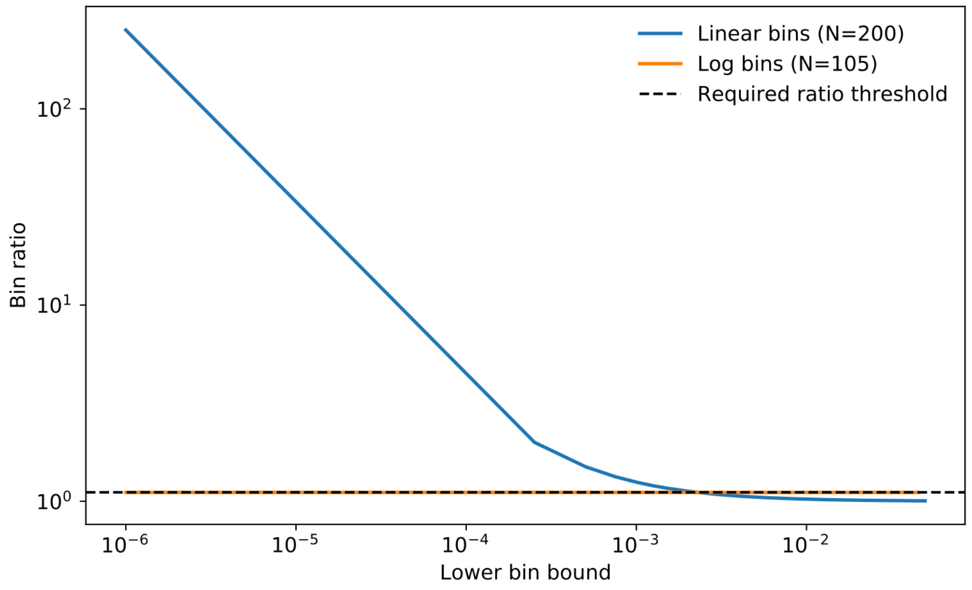


**Figure S3.** Comparison of bin-bound ratios for linear and logarithmic discretization. The ratio between consecutive bin bounds $u_i/l_i$ is shown across the feasible concentration range for linear (blue) and logarithmic (orange) discretization schemes. The dashed line denotes the maximum allowable ratio $\varepsilon/(\varepsilon - m_a)$ required to guarantee the specified linearization accuracy. Linear binning produces large ratios in the lowest concentration bins, violating this condition, whereas logarithmic binning maintains a constant ratio across bins and satisfies the constraint with fewer bins.

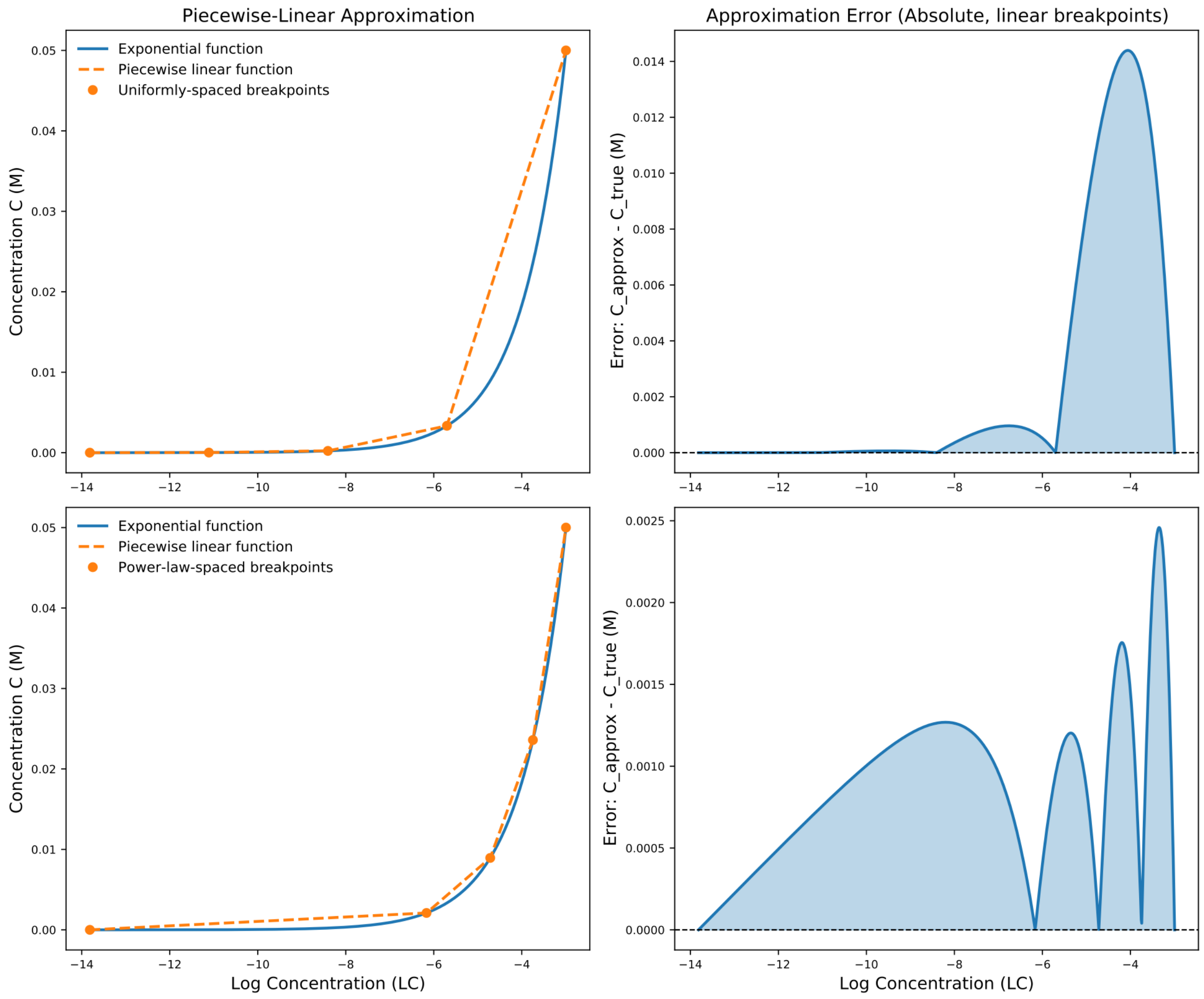


**Figure S4.** Impact of breakpoint distribution on the accuracy of the exponential linearization. The exponential function is approximated using piecewise-linear interpolation. The top row uses uniformly spaced breakpoints in log space, while the bottom row uses power-law-spaced breakpoints that concentrate points near the upper bound. The right panels show the corresponding absolute approximation error. Because the exponential function is strongly curved near the upper bound, uniform spacing produces large approximation errors in this region, whereas concentrating breakpoints there significantly reduces the error.

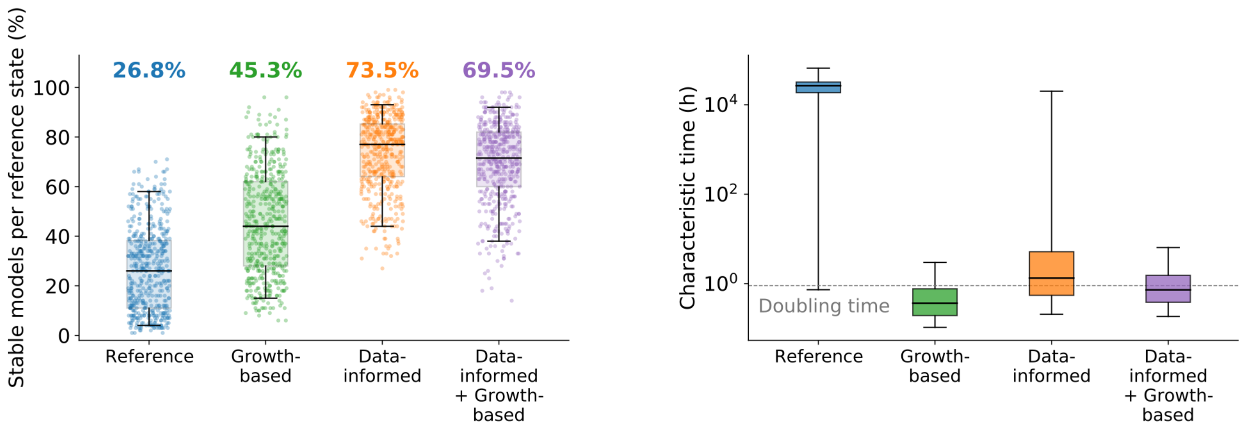


**Figure S5.** Combining data-informed and growth-based turnover constraints alters kinetic ensemble stability and characteristic times. The combined model applies literature-derived bounds to 18 metabolites and growth-based lower bounds to the remaining metabolites represented in the kinetic model. Left: Stable fractions among kinetic parameter sets per reference state and model category. Dots denote reference states, and labels denote overall stable fractions. Right: Characteristic times for stable parameter sets; the dashed line marks the cell doubling time. Boxes show medians and interquartile ranges, and whiskers span the 5th-95th percentiles.

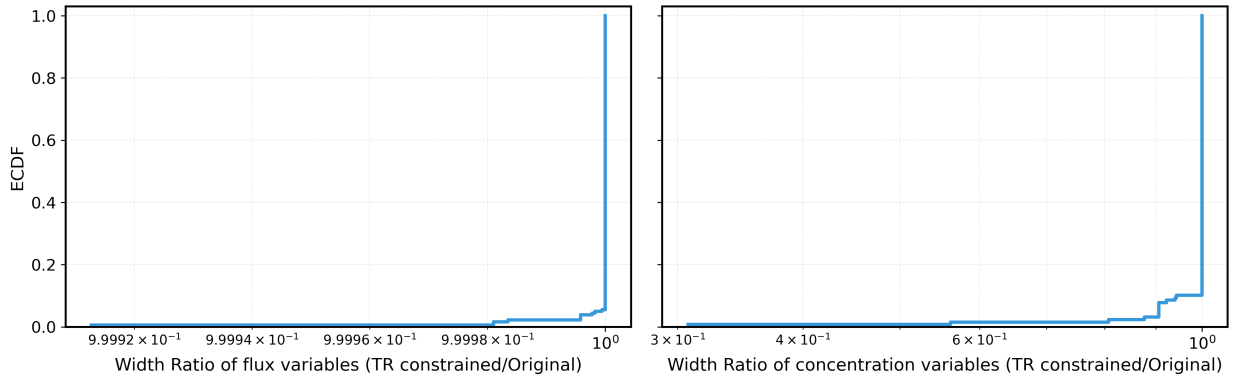


**Figure S6.** Effect of turnover-rate constraints on feasible ranges in the anaerobic *E. coli* model. Empirical cumulative distributions show the ratios of flux (left) and log-concentration (right) range widths under the growth-based 10× turnover cutoff relative to the original model. Ratios below 1 indicate narrower ranges. Flux ranges remained essentially unchanged, whereas a subset of log-concentration ranges narrowed.

**Table S1.** Distributional differences in reaction fluxes between steady states sampled with and without TR constraints. Values show the median Jensen-Shannon (JS) divergence and Wasserstein distance of flux distributions aggregated by metabolic subsystem. Larger values indicate greater shifts in the distribution of fluxes between the two sampling populations.

| Subsystem | Median JS divergence | Median Wasserstein distance |
|---:|---:|---:|
| Pyruvate Metabolism | 0.258 | $1.44\times10^{-1}$ |
| Glycolysis/Gluconeogenesis | 0.111 | $1.23\times10^{-1}$ |
| Pentose Phosphate Pathway | 0.015 | $2.47\times10^{-2}$ |
| ETC_Rxns | 0.101 | $1.71\times10^{-2}$ |
| Folate Metabolism | 0.028 | $1.43\times10^{-2}$ |
| Alternate Carbon Metabolism | 0.170 | $1.26\times10^{-2}$ |
| Transport, Inner Membrane | 0.111 | $1.09\times10^{-2}$ |
| Lipopolysaccharide Biosynthesis / Recycling | 0.140 | $9.68\times10^{-3}$ |
| Nucleotide Salvage Pathway | 0.104 | $9.08\times10^{-3}$ |
| Anaplerotic Reactions | 0.038 | $7.45\times10^{-3}$ |
| Inorganic Ion Transport and Metabolism | 0.074 | $4.06\times10^{-3}$ |
| Glyoxylate Metabolism | 0.196 | $3.60\times10^{-3}$ |
| Glutamate Metabolism | 0.100 | $3.44\times10^{-3}$ |
| Unassigned | 0.069 | $1.28\times10^{-3}$ |
| Alanine and Aspartate Metabolism | 0.070 | $1.10\times10^{-3}$ |
| Citric Acid Cycle | 0.052 | $1.10\times10^{-3}$ |
| Purine and Pyrimidine Biosynthesis | 0.132 | $1.57\times10^{-4}$ |
| Threonine and Lysine Metabolism | 0.132 | $1.50\times10^{-4}$ |
| Glycine and Serine Metabolism | 0.090 | $1.31\times10^{-4}$ |
| Arginine and Proline Metabolism | 0.132 | $1.04\times10^{-4}$ |
| Cysteine Metabolism | 0.132 | $1.00\times10^{-4}$ |
| Methionine Metabolism | 0.132 | $6.22\times10^{-5}$ |
| Cofactor and Prosthetic Group Biosynthesis | 0.132 | $3.15\times10^{-7}$ |
| Exchange | 0.132 | $2.71\times10^{-7}$ |

**Table S2.** Literature-derived metabolite turnover-time ranges calculated as lower and upper turnover time bounds. G, Gerosa (2015); P, Park (2016); K, Kochanowski (2021); T, Taymaz-Nikerel (2009).

| Metabolite | Model identifier | Lower bound (s) | Upper bound (s) | Studies |
|---:|---:|---:|---:|---:|
| KDPG | 2ddg6p_c | 0.07582 | 0.1014 | G |
| 6-Phosphogluconate | 6pgc_c | 0.4523 | 15.82 | G, P, K, T |
| cis-Aconitate | acon_c | 0.01659 | 0.04229 | K |
| α-Ketoglutarate | akg_c | 0.5806 | 0.6194 | T |
| Citrate | cit_c | 1.722 | 2.201 | G |
| Dihydroxyacetone phosphate | dhap_c | 0.6834 | 3.147 | G, P, K |
| Fructose 6-phosphate | f6p_c | 1.074 | 1.326 | T |
| Fumarate | fum_c | 0.6682 | 0.7318 | T |
| Glucose 6-phosphate | g6p_c | 3.448 | 3.752 | T |
| Isocitrate | icit_c | 0.01664 | 0.02912 | G |
| L-Malate | mal__L_c | 2.711 | 2.889 | T |
| Phosphoenolpyruvate | pep_c | 2.616 | 2.784 | T |
| Pyruvate | pyr_c | 1.3 | 1.7 | T |
| Ribulose 5-phosphate | ru5p__D_c | 1.708 | 2.174 | G |
| Sedoheptulose 7-phosphate | s7p_c | 2.303 | 41.1 | G, P, K |
| Succinate | succ_c | 7.859 | 9.941 | T |
| Succinyl-CoA | succoa_c | 0.4733 | 1.313 | P |
| Xylulose 5-phosphate | xu5p__D_c | 2.878 | 4.317 | G |
